# Spatially Resolved Transcriptomic Analysis of the Germinating Barley Grain

**DOI:** 10.1101/2023.01.24.525109

**Authors:** Marta Peirats-Llobet, Changyu Yi, Lim Chee Liew, Oliver Berkowitz, Reena Narsai, Mathew G. Lewsey, James Whelan

## Abstract

Seeds, which provide a major source of calories for humans, are a unique stage of a flowering plant ‘s lifecycle. During seed germination the embryo reactivates rapidly and goes through major developmental transitions to become a seedling. This requires extensive and complex spatiotemporal coordination of cell and tissue activity. Existing gene expression profiling methods, such as laser capture microdissection followed by RNA-seq and single-cell RNA-seq, suffer from either low throughput or the loss of spatial information about the cells analysed. Spatial transcriptomics methods couple high throughput analysis of gene expression simultaneously with the ability to record the spatial location of each individual region analysed. We developed a spatial transcriptomics workflow for germinating barley grain to better understand the spatiotemporal control of gene expression within individual seed cell types. More than 14,000 genes were differentially regulated across 0, 1, 3, 6 and 24 hours after imbibition. This approach enabled us to observe that many functional categories displayed specific spatial expression patterns that could be resolved at a sub-tissue level. Individual aquaporin gene family members, important for water and ion transport, had specific spatial expression patterns over time, as well as genes related to cell wall modification, membrane transport and transcription factors. Using spatial autocorrelation algorithms, we were able to identify auxin transport genes that had increasingly focused expression within subdomains of the embryo over germination time, suggestive of a role in establishment of the embryo axis. Together, our data provides an unprecedented spatially resolved cellular map for barley grain germination and specific genes to target for functional genomics to define cellular restricted processes in tissues during germination. The data can be viewed at https://spatial.latrobe.edu.au/.

## 1 Introduction

Seeds provide 70% of human calorie intake and are essential for sustainable food security. This unique stage of the life cycle of flowering plants not only allows plants to optimise survival strategies, but has also formed the basis of settled human civilisation. Consequently, plants have been bred for seed yield and quality since the foundation of agriculture and the study of breeding continues intensively^1^. While the genome sequence of agricultural crops greatly accelerates efforts to breed for desirable traits, it is the dynamic expression of the genome in the millions of cells that defines these traits. Thus, while whole-organ or tissue-specific transcriptome data is valuable in fundamental research and applications, recent technologies enabling the interrogation of gene expression at single-cell resolution have revolutionised our ability to understand plant growth and function at the basic level of organisation, i.e. the cell^2^.

Several approaches to obtain single-cell transcriptomes have been developed, with microfluidics-based single-cell sequencing (scRNA-seq) currently of wide use due to the high-throughput nature of this technology. The first scRNA-seq studies in plants were applied to root tissues^3-5^ because of their spatially defined developmental profile along the longitudinal axis and the relative ease of isolating individual cells. scRNA transcriptomes are also emerging for other organs, showing that this approach can be successfully applied to a variety of tissues^6-9^.

It has been proposed that the plant science community come together around the goal of creating a complete plant cell atlas, which should incorporate transcriptomic information about all cells in an entire plant^10^. scRNA-seq is considered a foundational technology for this goal, but the technology has limitations that prevent it being a full solution for the plant cell atlas. Some cell populations or types may not be assayed in scRNA-seq experiments, due to the varying size of plant cells, different cell wall compositions, the necessity and difficulty of optimising protoplast or nuclei isolation methods for individual tissue types, the requirements for substantial amounts of input tissue and the relative rarity of certain cell types. Moreover, scRNA-seq requires the dissociation of tissues into individual cells for microfluidic handling. This discards crucial information on the spatial origin of those cells, which defines their identity and function. As a result, scRNA-seq data analysis reconstructs cell identities and defines cell developmental trajectories by relying on cell-type-specific marker genes and gene co-expression. These approaches are bioinformatically challenging and may introduces biases^4,11,12^.

Spatial transcriptomics is an emerging approach in plant biology that promises to overcome some of the limitations of scRNA-seq^13,14^. This approach can be applied to any plant tissue that can be sectioned and morphology maintained, which has occurred extensively for many crop plants. When sectioning is optimised, it requires small amounts of tissue, employs fixation to preserve a full snapshot at a particular time, and retains cell spatial information, which are all very attractive qualities. These indicate spatial transcriptomics could make a significant contribution to obtaining cell-specific transcriptomes in plants, complementing scRNA-seq approaches and opening an exciting era for plant biology.

Few spatial transcriptomic studies have been reported in plants, contrasting with mammalian systems where these technologies have been successfully applied to gain insights into development and disease progression^15,16^. The first plant spatial transcriptomic study used a 100 µm resolution array to demonstrate the applicability of the method on the *Arabidopsis thaliana* inflorescence meristem, *Populus tremula* leaf buds and *Picea abies* female cones ^17^. A second study that used a different technology, Stereo-seq, with a 220 nm resolution array detected almost 20,000 expressed genes. Here, subtle differences were revealed in the expression of genes associated with photosynthesis across the *Arabidopsis* leaf cells. However, the authors noted that some cells associated with vascular bundles were missing^18^. Most recently, a gene expression map of flower development in orchids was generated using the 10x Visium technology at 55 µm resolution^19^ and this technology was also used to confirm the laser microdissection description of the spatial integration of C4 and CAM photosynthetic metabolism^20^. However, as yet no spatiotemporal studies with a plant developmental process have been reported.

Here, we present a time-series spatial transcriptomic analysis of germinating barley grain at 55 µm resolution. We detected the expression of >14,000 genes over five to six tissue types, at 0, 1, 3, 6 and 24 hours after imbibition (HAI). We analysed and distinguished the discrete spatial and temporal transcriptional activity between barley grain tissues and between domains within tissues. This allowed us to analyse the distinct and subtle expression domains of individual gene family members, for example ion and auxin transporters, transcription and translation and relate them to specific spatial functions. This spatial and temporal resolution in gene expression will provide specific targets for grain improvements and promoters to drive precise expression for synthetic biology approaches to alter grain quality.

## Results

### Establishing a spatial transcriptomics workflow for barley grains

To obtain the transcriptional signature of barley grains from thinly sliced sections whilst preserving spatial information, we optimised spatial transcriptomics using Visium technology (10x Genomics) for germinating barley grains (0, 1, 3, 6 and 24 HAI) (Supplementary Fig. 1a). The grains were cut in half, snap-frozen in an isopentane bath and embedded in optimal cutting temperature to generate cryoblocks (Supplementary Fig. 1b). We prepared eight µm thick longitudinal sections of the barley grains, then mounted them onto the active sequencing areas of Visium Gene Expression slides (6.5 × 6.5 mm^2^, 10x Genomics, Supplementary Fig. 1b). These active areas contain ∼5,000 spot arrays with a diameter of 55 µm and a centre-to-centre distance of 100 µm (Supplementary Fig. 1c). The spots on these slides are coated with spot-specific positional barcodes attached to oligo (dT) primers and unique molecular identifiers (UMIs) used for locating, capturing and identifying the mRNAs, respectively (see methods and Supplementary Fig. 1). In total, we prepared and analysed 20 samples, i.e. four sections per grain for each of the five time points (Supplementary Fig. 1b). To assess whether the detected transcript signal was stringently confined under individual plant cells, we used the Visium Tissue Optimization kit (10x Genomics) with modifications to suit barley grains (see methods and Supplementary Fig. 2, adapted from^21^. To further determine the optimal conditions for the starchy barley grain, we compare the sequencing quality of a pre-permeabilization method described in ^21^ and the standard permeabilization method (10x Genomics) using sections from two different seeds at the 24 HAI time point. A similar number of genes (c. 13,000) was detected in all six of these sections (Supplementary Fig. 2e) leading to the use of the standard permeabilization method for all subsequent gene expression experiments given its faster processing time.

### Spatiotemporal gene expression dynamics during barley grain germination

We applied the optimised spatial transcriptomics workflow to study spatiotemporal gene expression dynamics during barley germination in an unbiased manner. We collected grains across a time-series of 0 (dry), 1, 3, 6 and 24 HAI and obtained four serial sections (per time point) along their longitudinal axis. Space Ranger (10x Genomics) was used for primary data processing and gene expression data overlaid upon grain section images using Loupe Browser. Among all sections, the minimum number of spots covered by tissue is 1171 and the highest number of tissue-covered spots is 1858. Within those tissue-covered spots, the mean number of reads per spot ranges from 57,397 to 103,993. The median UMI counts per spot varied between 69 to 1689 with median genes per spot between 57 and 1092 across the different sections. (Supplementary Fig. 3, Supplementary Table 1). Note that the embryo tissues show much higher UMI and gene numbers (Supplementary Fig. 4, Supplementary Fig. 5). Secondary analysis used Seurat and SCTransform to generate clusters across four sections for each time point with the uniform manifold approximation and projection (UMAP) algorithm for dimensionality reduction^22,23^. Resulting outputs for 24 HAI are shown in Fig. 1a-d and for all other time points in Supplementary Fig. 6, 7, 8 and 9 for 0 HAI, 1 HAI, 3 HAI and 6 HAI, respectively. A total of 11 clusters were identified at 24 HAI based on histological features and spatial transcriptomic data, with one cluster each for the coleorhiza, scutellum and aleurone, three for the embryo and five for the endosperm (Fig. 1a-b, Supplementary Fig. 10). The four sections showed high correlation with a Pearson correlation coefficient from 0.89 to 0.96 at 24 HAI (Fig. 1c-d), and the high correlation across sections were also observed at other time points (Supplementary Fig. 11). Furthermore, the four sections showed a similar proportion of number of spots (Supplementary Fig. 12) and a high correlation (Supplementary Fig. 13, 14, 15, 16, and 17) for clusters at each time point. These results again showed the high reproducibility across different sections. Marker genes for each tissue/cluster were identified by comparison to their expression in all other clusters at all time points (Supplementary Table 2).

**Figure 1.**
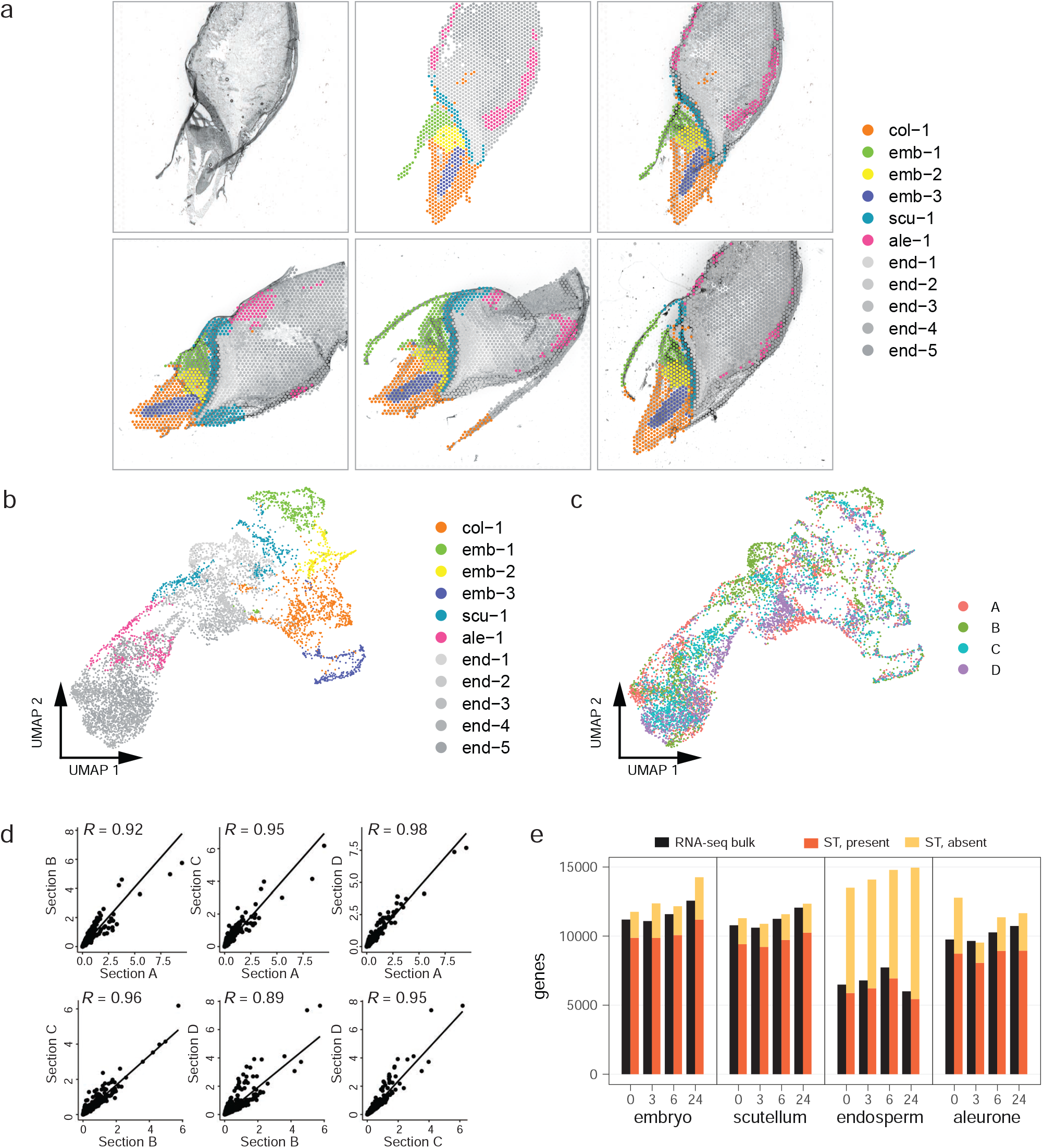
Spatially resolved transcriptome analysis of barley grain. **a**, Spatial visualization of the unbiased spot clustering for four 24 hours after imbibition (HAI) barley sections. Top left panel, bright field image of section D, top middle panel, spatial localization of each cluster of section D, top right panel, merged bright field image and spatial clusters of section D. Bottom panels, merged bright field image and spatial clusters of other three sections. The tissue/cell-type identity of each cluster was assigned based on the location of each cluster. Col: coleorhiza, emb: embryo, scu: scutellum, ale: aleurone, end: endosperm. **b**, Uniform manifold approximation and projection (UMAP) of spatial spots from four 24 HAI barley sections. Dots correspond to individual spots on the Visium slide; n = 6961 spots; colours indicate cluster association for each spot. **c**, UMAP of spatial spots from four 24 HAI barley sections. Colours indicate different sections for each spot. **d**, Pair-wise Pearson correlation between different sections for 24 HAI. Dots correspond to genes expressed in at least 5 spots. X axis and Y axis show the average of normalized count of tissue covered spots from two different sections, respectively. Pearson correlation coefficient is indicated as *R*. **e**, Number and overlap between genes detected by bulk RNA-seq of barley grain tissues and spatial transcriptomics. For genes identified by spatial transcriptomics, their presence or absence in the same location as in a tissue-specific bulk RNA-seq experiment^24^ was determined. The bar chart indicates the number of genes across comparable time points and tissues between both experiments. The cut-offs for inclusion in this comparison were at least five transcripts per million (TPM) for the RNA-seq experiment and at least five UMI per million for the spatial transcriptomics data, respectively.

We compared our spatial transcriptomic analysis to a published tissue-specific transcriptome analysis of barley grains generated by hand dissection to determine how closely the two approaches agreed with each other^24^. The tissue-specific analysis detected expression of 19,611 genes across all tissues, compared with 14,594 genes detected across our complete spatial transcriptomic dataset (Fig. 1e). Expression was next benchmarked per tissue. The spatial transcriptomic approach consistently detected between 83% and 90% of expressed genes present in the same tissue of the tissue-specific RNA-seq experiment, except for in the endosperm (Fig. 1e). Spatial transcriptomics detected twice as many expressed genes in the endosperm than tissue-specific RNA-seq, but with almost all the expressed genes detected by hand dissection also present in the spatial transcriptomic dataset (Fig. 1e). Note that the spatial transcriptomic data presented here comprises four sections of eight µm thickness each along the longitudinal axis of a grain that is around 5,000 µm wide. Therefore, in comparison to tissue-specific sequencing only a portion of the whole grain will have been sampled which might not include all cell types. Correlations of gene expression across the different time points between the embryo, scutellum and aleurone of the spatial transcriptomics and the tissue-specific RNA-seq were R^2^ ranging from 0.42 to 0.59. For the endosperm, no correlation was observed with R^2^ between 0.09 and 0.16 (Supplementary Fig.18). In bulk RNA sequencing the expression of a gene is averaged over thousands to millions of cells, and the expression can represent a high abundance in a few cells or uneven expression levels, which remains unresolved following Simpson ‘s paradox^25^. Based on these benchmarks, spatial transcriptomics on barley grain is comparable in depth to tissue-specific sequencing with the added advantage of spatial resolution that eliminates any potential errors that arise due to averaging of expression levels that may be present in a cellular landscape^26^.

Sets of expressed marker genes which were defined by comparing gene expression of different clusters pair-wisely (see methods) could be clearly defined for each cluster, which corresponded to individual tissues or regions within tissues, at 24 HAI (Fig. 2a). Again, we detected a higher number of UMIs and genes in the embryo tissues (Supplementary Fig. 19, Supplementary Fig. 20). In coleorhiza (cluster col-1) expression of a distinct set of four genes was observed, with a second set that overlapped the radicle (cluster emb-3) (Fig. 2a, Supplementary Table 3). The latter is not unexpected given the relationships between radicle and coleorhiza. The scutellum (cluster scu-1) expressed a distinct set of marker genes as well as some additional marker genes that overlapped with the endosperm, which is again not unexpected. The embryo was divided into three clusters. Of these, the coleoptile (cluster emb-1) expressed a distinct set of marker genes, whilst the radicle (cluster emb-3) and mesocotyl (cluster emb-2) expressed some overlapping marker genes that were differentiated by magnitude of expression. The endosperm was partitioned into five clusters, that could not be distinguished from one another by small sets of marker transcripts. It is therefore likely that the endosperm clusters arose from more complex or subtle differences in gene expression patterns. These five clusters may reflect differential water uptake and mobilisation of reserves across the endosperm upon imbibition, demonstrating the spatial heterogeneity in the magnitude of expression of genes between cells in the same tissue. Considered together, these results demonstrate the advantage of the spatial transcriptomic approach. It allows tissues and regions within tissues to be distinguished by a combination of their physical location and gene co-expression, rather than by co-expression alone. We were consequently able to manually assign and curate spot locations to tissues without solely relying on marker genes or bioinformatic algorithms.

**Figure 2.**
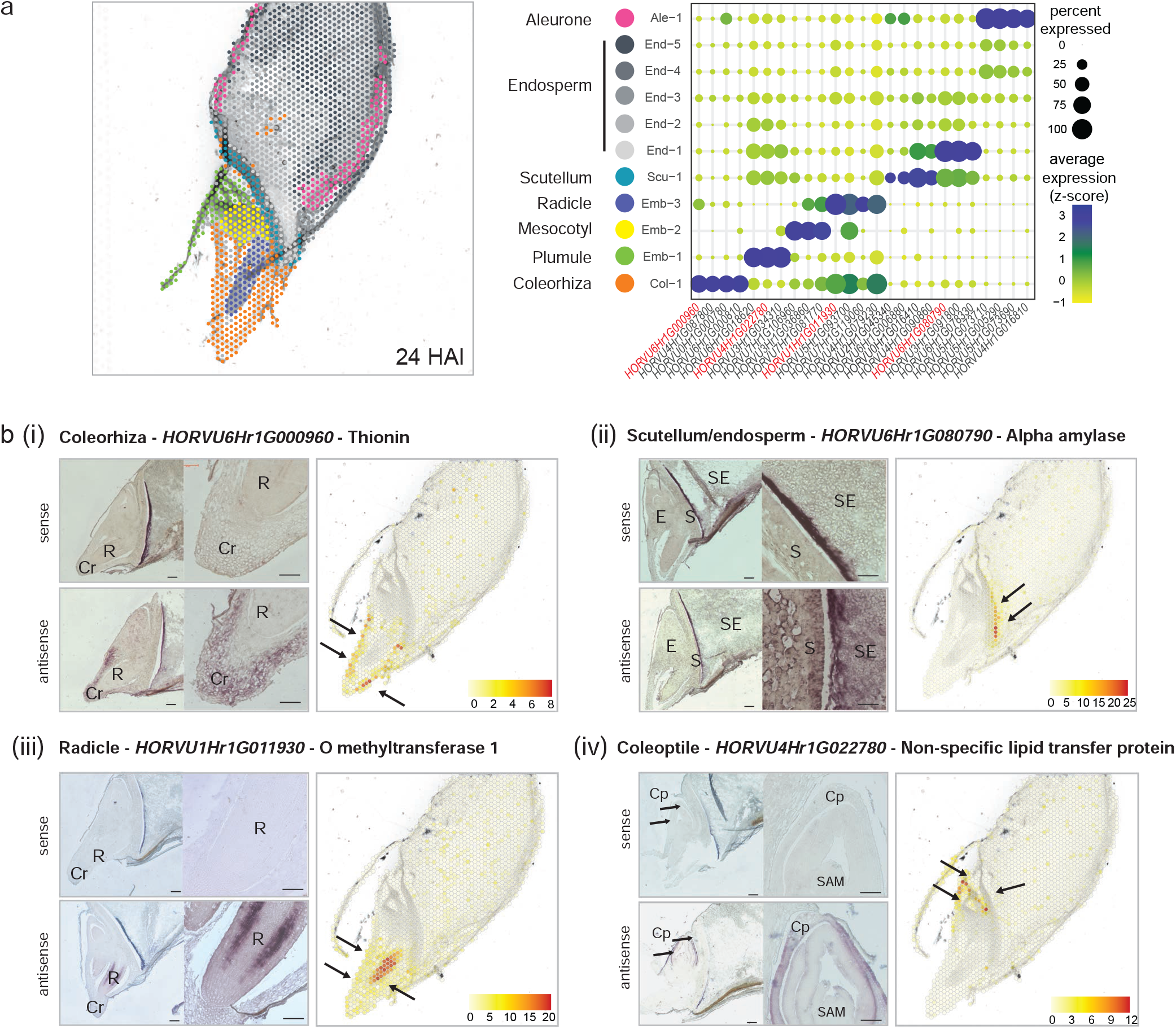
Identification of tissue-specific marker genes and their validation by *in situ* hybridisation. **a**, Spatial representation of the 11 clusters determined at the 24 HAI timepoint (left panel) and bubble plot (right panel) showing transcript enrichment (average expression and percentage) of representative cell type-specific marker genes in the 11 clusters. Spot colours correspond to the same tissues in the bubble plot. Genes highlighted in red were validated by *in situ* hybridisation. **b**, *in situ* hybridization of selected marker genes (i) *HORVU6Hr1G000960*; (ii) *HORVU6Hr1G080790*; (iii) *HORVU1Hr1G011930*; (iv) *HORVU4Hr1G022780* confirmed localization of tissue-type specific transcripts at 24 HAI timepoint. For each *in situ* hybridisation, top panels show hybridisation with the sense probes (negative controls) and bottom panels with the antisense probes. *In situ* images on the right correspond to 40x magnifications of images on the left. Scale bars correspond to 200 µm and colour scales represent normalised counts. Coleorhiza (Cr, Col), Radicle (R), Coleoptile (Cp), Shoot apical meristem (SAM), Aleurone (A, Ale), Starchy endosperm (SE, End), Embryo (E, Emb), Scutellum (S, Scu). Black arrows indicate the expression areas.

### Validation of spatial expression by RNA *in situ* hybridisation

The expression patterns of transcripts detected by spatial transcriptomics were orthogonally validated using RNA *in situ* hybridisation (Fig. 2b). *THIONIN* genes encode low molecular weight basic cysteine-rich antimicrobial peptides and are highly expressed early in germination in a tissue-specific manner in rice coleoptiles^27^. Spatial transcriptomic analysis at 24 HAI indicated a coleorhiza-specific expression pattern for a *THIONIN* gene (*HORVU6Hr1G000960*) (Fig. 2b i). This was confirmed by RNA *in situ* hybridisation (Fig. 2b i). α-amylases are critical enzymes for the mobilisation of starch reserves in germinating barley grains and have a scutellum and aleurone expression pattern^28^. Spatial transcriptomics detected *HORVU6Hr1G080790* gene expression in the scutellum at 24 HAI, with strongest expression at the ventral end (Fig. 2b ii). RNA *in situ* hybridisation confirmed a scutellum expression pattern but could not distinguish expression magnitude across the scutellum. Any localised gradient in expression is difficult to resolve by RNA *in situ* hybridisation due to the non-linear nature of colorimetric detection methods that are end-product inhibited. For the radicle, the specific expression of an *O METHYLTRANSFERASE 1* gene (*HORVU1Hr1G011930*) involved in lignin biosynthesis was confirmed^29^ (Fig. 2b iii). Coleoptile and leaf specific expression of a gene encoding a non-specific lipid transfer protein (*HORVU4Hr1G022780*) has been previously reported in barley^30,31^. A similar pattern was observed here by RNA *in situ* hybridization and spatial transcriptomics (Fig. 2b iv).

Thus, the spatial transcriptomics results for germinating barley grain were consistent with the localisation, abundance and depth obtained with other approaches, but had the advantage of combining all data into a single high-throughput approach.

### Conservation of spatial expression patterns across species

Given the extensive transcript profiling studies in *Arabidopsis* tissues we undertook an orthology approach to determine if the marker genes defined in this study also displayed tissue specific expression in *Arabidopsis*. For the top 10 genes per cluster based on specificity, per time point the *Arabidopsis* orthologs were defined using OMA^32^. This identified 14 *Arabidopsis* genes for endosperm, 25 genes for aleurone, 44 genes for scutellum and 145 genes for embryo. Note that most genes have a many-to-many relationship in each orthogroup, however paralogs were not included in this number. Analysis of the tissue expression pattern for these genes using Genevestigator^33^ revealed that only 10% to 30% of the marker genes defined in barley displayed a restricted expression in a comparable tissue in *Arabidopsis* (Supplementary Fig. 21, 22, 23, 24 and 25). Thus, the spatial expression of genes in seeds of the two species shows limited conservation.

### Spatiotemporal analysis of key biological processes during germination of barley grains

To reveal spatial and temporal patterns in the function of expressed genes, functional categories present in at least three time points per tissue or containing at least five genes at any time point were visualised, as well as categories present across the same time point in at least two tissues (Fig. 3a). The number of genes in the 26 identified functional categories are indicated as black data bars, and this revealed the spatiotemporal patterns linking expression and function (Fig. 3a). Four major functional groups were identified including DNA/RNA metabolism and translation, seed storage and metabolism, transport and miscellaneous functions (Fig. 3a). The majority of genes encoding transcription and translation functions were expressed in the embryo. Specifically, 56 to 133 genes encoding ribosomal proteins were expressed from 0 HAI to 24 HAI in the embryo tissue, peaking at 24 HAI. A temporal pattern for these was also observed with the expression of 51 and 89 genes encoding ribosomal proteins highly expressed at 24 HAI in the scutellum and endosperm, contrasting with much fewer, if any, observed at other time points for these tissues (Fig. 3a). A similar pattern of expression was also observed for the genes encoding transcription factors, elongation factors and heat shock factors, indicating an embryo specific expression pattern, peaking at 24 HAI for many of these genes (Fig. 3a). Notably, functions that were associated with translation, ribosomal proteins, elongation factors and heat shock proteins (chaperones), were expressed in the embryo at all time points, compared to other tissues. This spatial gene expression pattern suggests that the ability of germination to proceed without transcription, but requiring translation^34,35^, is an embryo specific function.

**Figure 3.**
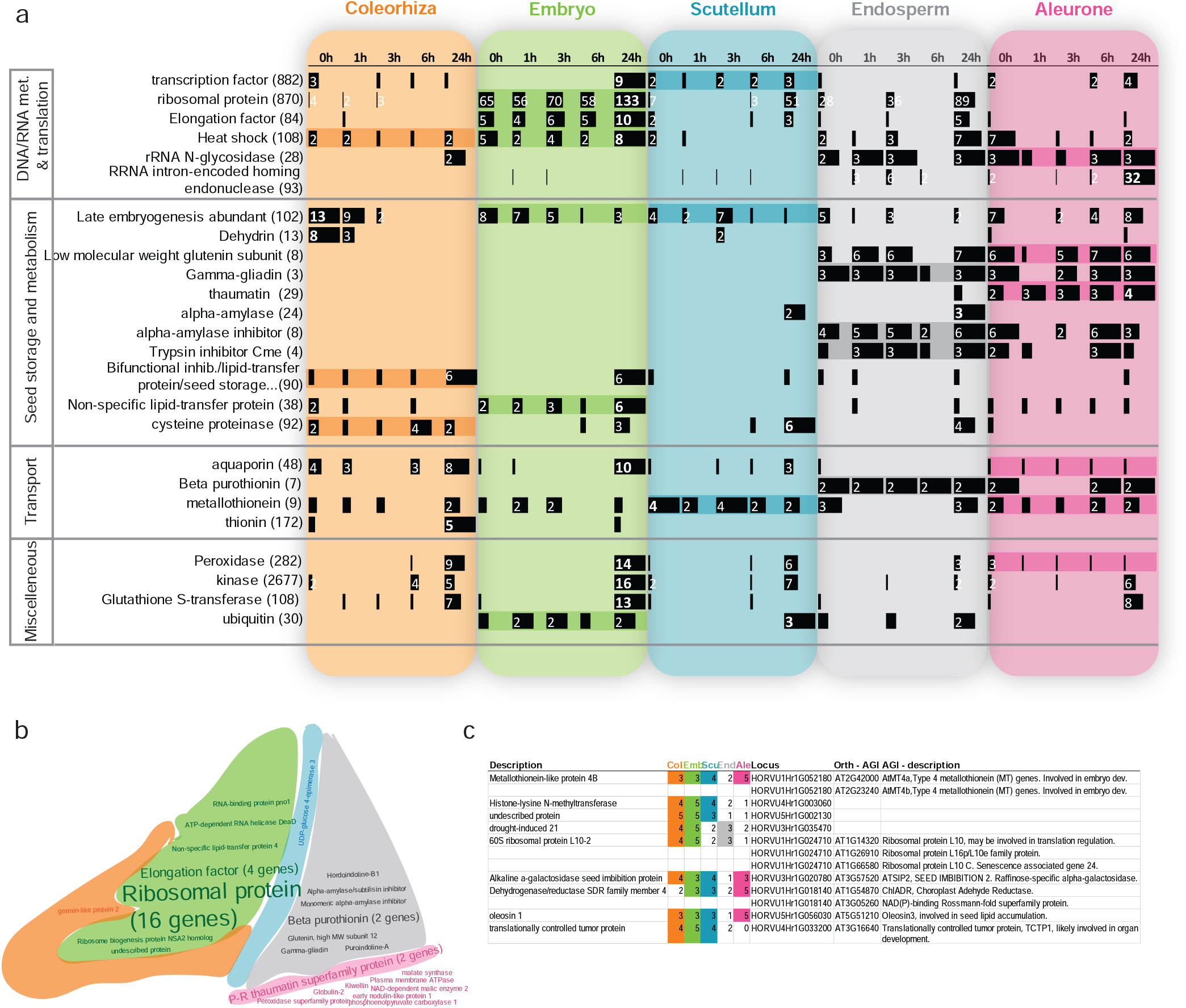
Spatiotemporal analysis of key biological processes during barley germination. **a**, Tissue specific functional category analysis. The categories identified from wordcloud outputs for the gene lists at each time point and tissue are shown, for those categories identified in at least three time points for a given tissue type or those with >5 genes at a given time point. The numbers of genes expressed at each time point and tissue are shown for each functional category using black databars across the rows. The white numbers indicate the genes corresponding to that category at the specific tissue and time point. Bars with no number correspond to 1. Bold white numbers on the bars indicate the category maximum number. The number of genes in the genome for that category is indicated in parentheses. **b**, Wordcloud categories representing highly tissue specific genes. There were 45 genes identified as having highly tissue specific expression. These were highly expressed in all 5 time points for one tissue, with no expression or specific expression in just one time point in other tissues. **c**, Most highly expressed genes across all samples. The table shows the genes expressed in 15 or more of the 25 samples making up all time-points and tissues. The number of time points at which a gene is expressed for each tissue is indicated in coloured columns as well as the Arabidopsis orthologue(s) and its description.

Distinct functions were also expressed in the endosperm and aleurone. Genes encoding rRNA N-glycosidase and rRNA intron-encoded homing endonucleases showed more specific expression in the aleurone and endosperm, with 32 of the 93 genes in the genome encoding the latter showing highly specific expression at 24 HAI in the aleurone (Fig. 3a). Similarly, genes encoding seed storage and metabolism functions also showed more specific expression in the endosperm and aleurone. Notably, genes encoding late embryogenesis abundant (LEA) proteins also showed temporal specific expression with the highest number of genes expressed at 0 HAI in four of the five tissue types, with the highest number (13 genes) expressed in the 0 HAI coleorhiza tissue (Fig. 3a). Dehydrins, which are a type of LEA that are known to have important roles associated with phenotypic traits^36^ also showed a similar pattern with the eight of the 13 dehydrin encoding genes in the genome expressed at 0 HAI in the coleorhiza (Fig. 3a). Based on studies in wheat^37^, the expression of prolamin encoding genes such as the low molecular weight glutenin and gamma gliadin genes were highly specific to the endosperm and here we see this specificity in the aleurone as well (Fig. 3a). This pattern was also observed for genes encoding seed metabolism functions, including alpha-amylase, alpha-amylase inhibitor and trypsin inhibitor CME, which is also known to be specific to the endosperm in barley^38^. Four genes encoding pathogenesis-related thaumatin like proteins were expressed specifically in the aleurone, with thaumatin like proteins known to have roles in defence, development and seed germination with specific expression observed during germination^39^. In contrast to starch metabolism, lipid metabolism functions including bifunctional inhibitor/lipid transfer protein as well as non-specific lipid transfer protein encoding genes showed more specific expression in the coleorhiza and embryo specifically, with the peak number of genes expressed at 24 HAI (Fig. 3a).

Aquaporin and thionin encoding genes were expressed specifically in the coleorhiza and embryo, peaking at 24 HAI. Aquaporins are important proteins involved in the movement of water in the seed^40^ and expression of aquaporin genes was seen from 0 HAI in the coleorhiza, while thionin encoding genes were only observed at 24 HAI indicating temporal specific regulation in the expression of these genes as well (Fig. 3a). Notably, of the seven genes encoding beta purothionins, which are involved in defence (as are thionins) and are known to be expressed in wheat endosperm^41^, two genes were expressed in all five time points in the endosperm and three time points in the aleurone, indicating highly specific expression (Fig. 3a). It has been shown that beta-purothionin can modify lipid packing and form channels at the bilayer surface to increase water accessibility in the interfacial region^41^, thus the genes showing these specific expressions are worth exploring further. Interestingly, four genes encoding metallothionins, from the nine genes encoding these in the genome, showed highly specific expression in the scutellum (Fig. 3a), collectively indicating that defence and water transport genes are tightly regulated in a spatio-temporal specific manner during seed germination in barley. Lastly, for the genes encoding peroxidases, kinases and glutathione-s-transferases, it is evidenced that these show a more temporal specific expression pattern with the greatest number of these expressed at 24 HAI and more specifically in the embryo (Fig. 3a).

We also identified 45 genes showing exclusive spatial specific expression patterns, defined by expression in all five time points in a particular tissue, with no specific expression or expression in just one time point in other tissues (Fig. 3b). In this way, 16 genes encoding ribosomal proteins, four genes encoding elongation factors, other three other genes with RNA binding functions were revealed embryo specific in expression (Fig. 3b). Similarly, a gene encoding UDP-glucose 4-epimerase 3 was scutellum specific in expression (Fig. 3b), and is orthologous to AtUGE1 and AtUGE3, which are known to be co-regulated with carbohydrate catabolic enzymes and have roles in growth and pollen development^42^. Whilst no endosperm specific genes met the above criteria for specificity, eight genes were identified which were expressed at all five time points in the endosperm and at least two time points in the aleurone, shown near the endosperm-aleurone junction (Fig. 3b, Supplementary Table 4). Notably, two genes encoding beta-purothionin were identified as endosperm-aleurone specific (Fig. 3b) and these are known endosperm specific potent antimicrobial proteins in wheat^43^. Similarly, gamma-gliadin and glutenin encoding genes were identified as endosperm-aleurone specific here (Fig. 3b), and these are known endosperm storage proteins in wheat, with significant roles in affecting flour quality^44^. Interestingly, two pathogenesis-related (P-R) thaumatin superfamily protein encoding genes showed aleurone specific expression across all time points (Fig. 3b), whilst another two were also aleurone specific but at 24 HAI only (Fig. 3a). The Arabidopsis orthologue to one of the two aleurone specific genes (*HORVU4Hr1G002650*) is osmotin 34 (AtOSM34), which has been shown to function as a positive regulator of ABA responses is under post-translational control^45^, thus further investigation into the function of these in barley germination could reveal insight into whether this regulatory role is conserved.

Lastly, to identify genes that were not specific to a given tissue, genes that were expressed in at least 15 of the 25 samples making up the five time-points in the five tissues were examined, revealing nine genes (Fig. 3c). Six of these had 10 *Arabidopsis* orthologues (OMA browser^32^), all of which were expressed across the developmental tissues (BAR). Apart from *At3G57520* and *At3G16640*, which showed moderately high expression during germination, the other eight genes showed the highest expression in dry seeds and/or during seed germination. For example, all three *Arabidopsis* genes that were orthologous to the barley 60S ribosomal protein L10-2 (*HORVU1Hr1G024710*) were co-expressed showing maximum expression in the germinating seed specifically. Similarly, both *Arabidopsis* genes orthologous to barley metallothionein-like protein 4B (*HORVU1Hr1G052180*) and both *Arabidopsis* genes orthologous to dehydrogenase/reductase SDR family member 4 (*HORVU1Hr1G018140*) showed maximal expression in the dry seed, just prior to germination. Notably, reducing expression of both *AtMT4a* and *AtMT4b* resulted in reduced seed weight and affected post-germinative early seedling growth, with over-expression resulting in the opposite effects^46^. The same study also revealed these to have a role in zinc accumulation in seeds, thus this gene in barley could represent a conserved orthologue, possibly having a similar role. Likewise, the other *Arabidopsis* genes that were orthologous to other genes in barley (Fig. 3c) have been shown to have roles in seed lipid accumulation, embryo development and organ development (Fig. 3c), thus the conserved expression revealed here could suggest these genes may have similar functional roles in seeds and development in barley. Interestingly, the three genes encoding a histone-lysine N-methyltransferase (*HORVU4Hr1G003060*), an undescribed protein (*HORVU5Hr1G002130*) and drought-induced 21 (*HORVU3Hr1G035470*) did not have significant orthologues in Arabidopsis. Considering their expression in all five tissues and at multiple time points for most tissues suggest these could be excellent targets to examine for functional roles in seed germination in barley.

### Aquaporin gene family members have distinct spatiotemporal expression patterns

To dissect further the spatial expression of genes, we examined in greater resolution the expression of some gene families associated with the functions outlined above. Transport of water, ions, solutes and other elements is essential for successful germination and occurs during imbibition in the earliest phase of germination^47^. We assembled a list of genes encoding proteins involved in transport processes, focusing on aquaporins (Fig. 4a, full list in Supplementary Fig. 26). Aquaporins are best known for their ability to facilitate water flow, but also transport other substrates such as various ions, solutes and CO ^48^. They are located in the plasma membrane (plasma membrane intrinsic proteins, PIPs) or the tonoplast (tonoplast intrinsic proteins, TIPs). The expression patterns of several TIPs and PIPs were assessed (Fig. 4a). Interestingly, at 24 HAI, we found highly expressed isoforms in the mesocotyl, scutellum and coleorhiza (Fig. 4b), illustrating that expression of aquaporin family genes is highly spatially distributed and organised during germination. The *HORVU4Hr1G079230* gene encodes a tonoplast intrinsic protein 1;1 (TIP1;1) that is the most abundantly expressed isoform of the TIP family^49^, which was also confirmed by our data. TIP1;1 is a housekeeping aquaporin and establishes a basal level of tonoplast hydraulic conductance^50,51^. In the spatial transcriptomic data, it was expressed mainly in the coleorhiza at early timepoints (0-6 HAI), then expression strongly increased at 24 HAI in coleorhiza and in the radicle and mesocotyl of the embryo (Fig. 4c, top). Another aquaporin gene, *HORVU1Hr1G043890*, encodes TONOPLAST INTRINSIC PROTEIN 3;1 (HvTIP3;1) which accumulates in the aleurone and outer layers of the barley seed^52^. We found this gene to be highly expressed in the embryo tissues at 0 and 1 HAI, followed by expression in the scutellum and aleurone layer at 3 and 6 HAI and in some areas of the aleurone/subaleurone layers at 24 HAI (Fig. 4c, bottom). These results illustrate the different spatiotemporal expression patterns of gene family members and how specific aquaporins regulate water transport in distinct tissues during seed germination.

**Figure 4.**
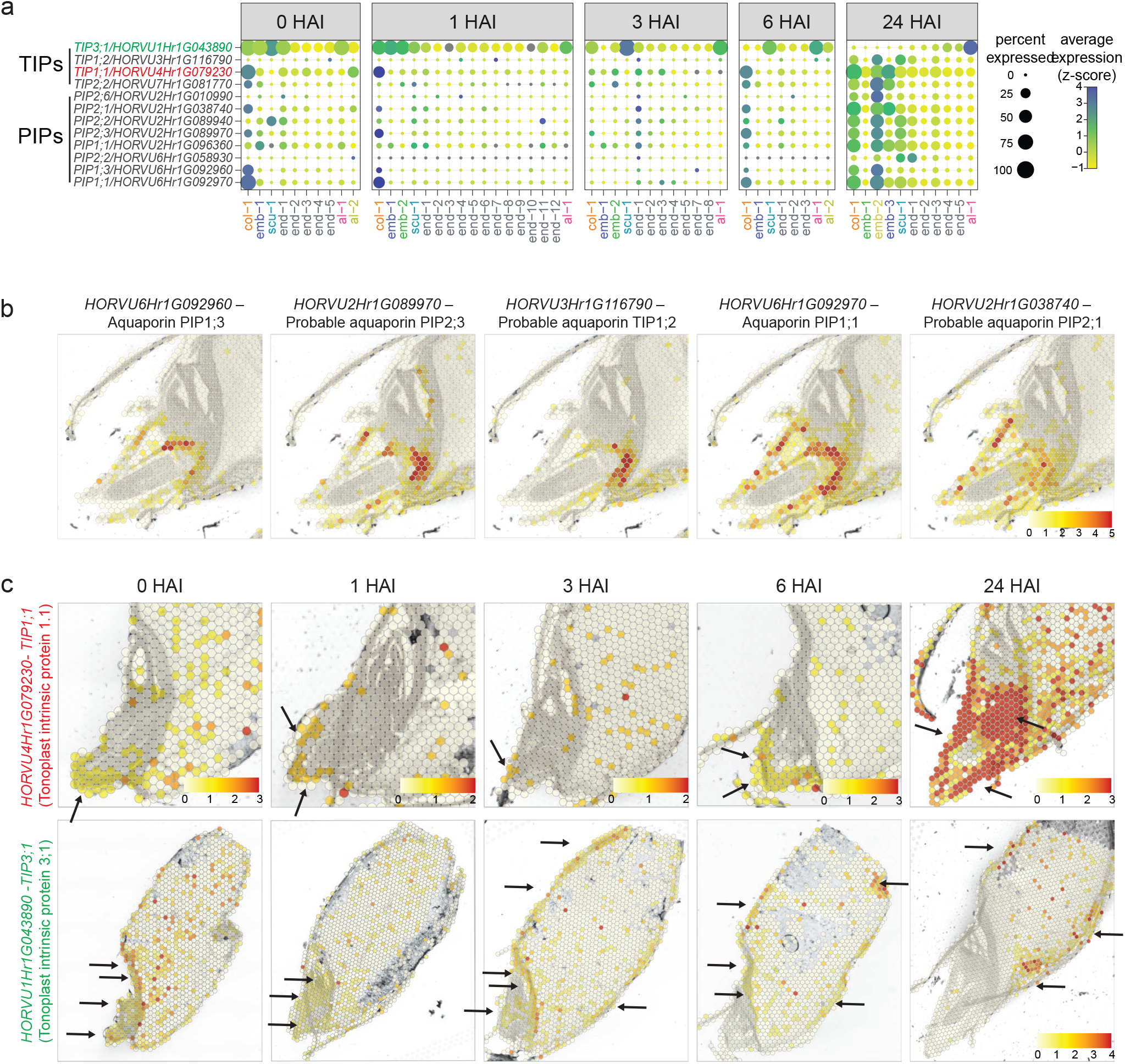
Spatio-temporal expression of aquaporin genes during barley germination. **a**, Bubble plots showing the spatially resolved expression of aquaporin gene family members (plasma membrane intrinsic proteins, PIPs; tonoplast intrinsic proteins, TIPs) across the experimental time-course. Circle size indicates the percentage of expression and colour indicates average expression after z-scoring. The x-axis represents individual clusters identified across tissues. For genes highlighted in green and red expression patterns are also shown in c). **b**, Highly expressed aquaporin family genes and their specific locations at 24 HAI. **c**, Time course expression of TIP genes: *HORVU4Hr1G079230* (TIP1;1) gene at the top panel and *HORVU1Hr1G043890* gene (TIP3;1) at the bottom panel. Colour scales represent the normalised counts with the same cut-off in each row or with cut-offs individually specified on the image. Black arrows indicate areas of expression. col: coleorhiza, emb: embryo, scu: scutellum, ale: aleurone, end: endosperm.

### Expression dynamics of cell wall metabolism genes

Transcription of genes involved in cell wall metabolism are up-regulated during the early stages of seed germination^53^ (Supplementary Fig. 27). Cell wall degradation, modification and precursor synthesis are essential processes in weakening of the tissues, e.g during radicle expansion. We detected spatially distinct and dynamic expression of cell wall-related genes across barley grain germination. For example, *HORVU3Hr1G073780*, encoding a type of cellulose synthase, was expressed at a relatively low level up until 6 HAI, but increased at 24 HAI (Supplementary Fig. 28a). It further displayed a very particular spatial pattern in the radicle, possibly the epidermis and cortex, suggesting this gene might be involved in new cell wall formation. Expansins were originally identified because of their ability to induce cell wall extension and are key regulators of cell growth^54^. We found that *ALPHA EXPANSIN 2* (*HORVU3HR1g081180*) was highly expressed in the late germination phase (6 and 24 HAI) around the expansion zone of the radicle (Supplementary Fig. 28b), implicating it in radicle protrusion for germination. Another cell wall-related gene, *HORVU5HR1g080860*, encodes a UDP-glucose epimerase (*HvUGE*) which participates in sugar precursor synthesis for cell wall biogenesis. Despite being identified in barley developing seed, leaf tips and mature roots^55^, our spatial study located *HvUGE* in the scutellum at all time points of germination (Supplementary Fig. 28c), which suggests this epimerase participates in cell wall polysaccharide synthesis during several barley developmental stages.

### Transcription factors involved in the spatiotemporal gene regulatory landscape

We sought insight into the gene regulatory landscape in barley by examining the expression of genes encoding transcription factors (Supplementary Fig. 29). One of the strongest expressed genes encoding a transcription factor was *HORVU1Hr1G086580*, a bZIP type transcription factor that responds to karrikin. Karrikins are small organic compounds known to be promoters of germination^56^. The spatial expression of *HORVU1Hr1G086580* is highest in early germination (0-3 HAI) in the scutellum and the adjacent endosperm area and then declines during germination (Supplementary Fig. 30b i). An *Arabidopsis* ortholog, *DELAY OF GERMINATION 1-LIKE 4 (DOGL4/AT4G18650)*, is a maternally expressed imprinted gene found in the endosperm of dry seeds and was described as a negative regulator of seed dormancy^57^. The *HORVU4Hr1G009730* gene encodes the *BARLEY KNOX3 (BKN3)* transcription factor that plays a pivotal role in shoot apical meristem (SAM) formation and maintenance as well as some morphogenetic processes during plant development^58^. We detected no expression of *BKN3* during early phases of germination (0-6HAI), but a very distinctive and localised expression at 24 HAI in the SAM (Supplementary Fig. 30b ii). The *HORVU2Hr1G104040* gene encodes a novel myc-related bHLH transcription factor and its ortholog in *Arabidopsis, Phytochrome-interacting factor 1 (PIF1)*, mediates light-regulated control of seed germination^59,60^. Our analysis showed that it is not expressed during early stages of germination (0-6 HAI) but expressed at 24HAI in the scutellum (Supplementary Fig. 26b iii). Another transcription factor, *HORVU7Hr1G026940*, encodes a DREB subfamily A-5 of ERF/AP2 transcription factor. DREBs are dehydration response element binding factors and one of the main groups of transcription factors that regulate expression of abiotic stress-inducible genes^61^. Spatial transcriptomics showed the expression of this transcription factor was restricted to the scutellum and endosperm during the early time points of germination (0-3 HAI) but it increased and moved to the aleurone layer at 6 HAI and 24 HAI (Supplementary Fig. 30b iv).

It is clear from the above examples for critical processes in germination that considerable variation in transcript abundance occurs with tissues (i.e. clusters) that has previously gone undetected.

### Non-biased discovery approaches define spatiotemporal domains during seed germination of barley

Expression of genes is not necessarily uniform across all cells within a tissue or cluster. Non-uniform expression within tissues reflects spatial organisation of function within that tissue. We assessed the ability of spatial transcriptomics to identify genes that had discrete patterns of spatial expression within tissues. To do so we focussed on the embryo tissues across all time points and applied Moran ‘s I spatial autocorrelation statistics to all expressed genes. This metric defines the correlation of gene expression of a local spot with its neighbouring spots and thus is a measure for non-random spatial patterns (see method for details)^62^. Moran ‘s I varies from -1 to 1, and a positive value indicates that cells with similar gene expression patterns are clustered together in a coherent domain. A value close to zero designates random spatial distribution of similar gene expression patterns, whilst a negative value shows that cells with similar gene expression patterns are spatially separated from each other. We considered genes that had a Moran ‘s I > 0.25, an adjusted p value < 0.01, expression in more than 5% of spots in a section, and meeting those thresholds in at least two sections for a time point, as having coherent expression domains; these are subsequently referred to as spatially variable genes (SVGs)^63^. The lowest number of SVGs was detected at 3 HAI and the highest number at 24 HAI (Fig. 5a, Supplementary Table 5). Among those SVGs, there were 204, 147, 68, 201 and 1098 SVGs consistently detected in all four sections for 0 HAI, 1 HAI, 3 HAI, 6 HAI, and 24 HAI, respectively (Supplementary Fig.3, Supplementary Table 5).

**Figure 5.**
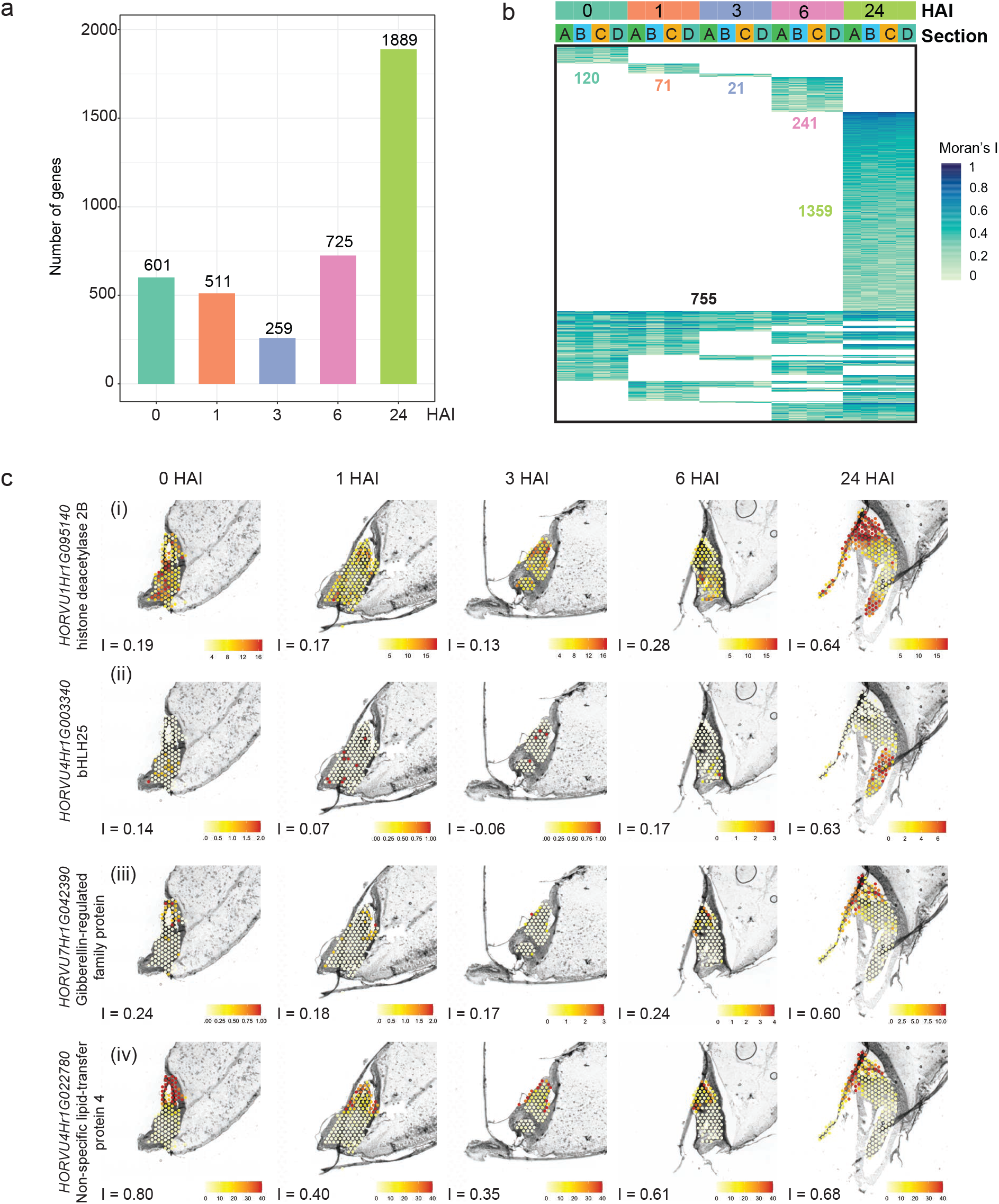
Spatial autocorrelation defines domains in the embryo of the germinating barley grain. **a**, Number of SVGs showing significant spatial coherence for each time point. For each section, Moran ‘s I was calculated for embryo-expressed genes, and genes with Moran ‘s I > 0.25, adjusted p < 0.01 and expressed percentage in embryo spots > 5% were considered having significant spatial coherence within a domain of the embryo. For each time point, SVGs showing significant spatial heterogeneity in at least two sections were plotted. **b**, Heatmap showing the Moran ‘s I of 2567 SVGs showing significant spatial coherence. The coloured text and black text indicate the number of time point-specific and time point-common genes, respectively. **c**, Examples of SVGs increasing spatial coherence. Colour scales represent normalised counts. I, Moran ‘s I value.

Many SVGs also had temporally specific expression; 1,812 SVGs were specific to one time point whilst 755 were spatially variable in more than one time point (Fig. 5b). Combined spatiotemporal expression patterns of SVGs were diverse. For example, a histone deacetylase 2B (HD2B, Fig. 5c i) with a Moran ‘s I = 0.64 at 24 HAI was enriched at the active growth centres including the radicle and shoot meristem in the embryo. HD2B histone deacetylase is associated with seed dormancy in *Arabidopsis*^64^. Two components of the histone deacetylation complex positively regulate seed dormancy while inhibiting seed germination by integrating ABA, ethylene and auxin signalling pathways^65,66^. *HORVU4Hr1G003340*, an ortholog of *Arabidopsis* bHLH25, is a transcription factor enriched in the radicle at 24 HAI (Moran ‘s I = 0.63, Fig. 5c ii). Overexpression of *Arabidopsis* bHLH25 results in increased lateral root numbers^67^. *HORVU7Hr1G042390* encodes a homolog of the GAST1 PROTEIN HOMOLOG 4 (GASA4) protein, which is a regulator of gibberellic acid-dependent seed germination and expressed in the shoot apex and embryo in *Arabidopsis*^68,69^. Its expression was focussed on the coleoptile with Moran ‘s I = 0.6 (Fig. 5c iii), consistent with the important role of GA in seed germination^70,71^. Moreover, the previously identified coleoptile-specific gene, *HORVU4Hr1G022780*, encoding a NON-SPECIFIC LIPID-TRANSFER PROTEIN 4, was also identified as an SVG (Fig. 5c iv).

Groups of SVGs may be expressed within similar spatial patterns because they have related or associated functions in the underlying cells. To examine this potential spatial co-expression we performed a high dimensional co-expression network analysis with the 1,889 significant SVGs from 24 HAI sample using hdWGCNA^72^. We determined seven co-expression modules and visualized their spatial expression patterns using eigengenes, which represent the common expression pattern of all genes within a module (Fig. 6a, Supplementary Table 6). Gene functions within modules were assessed using gene ontology (GO) enrichment analysis (Fig. 6b). Genes in the green module were highly expressed in the apex of radicle and were enriched for functions in glutathione (GSH) metabolic process and sulfur compound metabolic process. GSH is critical for (embryonic) root development in *Arabidopsis*, partly via modulation of auxin transport and signalling^73-75^. The turquoise module was expressed most highly at the base of the radicle and enriched for cell wall biogenesis and organization. Plant cell wall biogenesis and organization are important developmental processes for radicle emergence^76-78^. The brown module was expressed most highly in the mesocotyl and genes within this module were associated with water and fluid transport, which corresponds to the presence of the vascular tissues in this region. The yellow module was expressed most highly in the coleoptile and consisted of genes enriched in photosynthesis and chlorophyll biosynthetic processes. This corresponds to the role of the coleoptile as the first photosynthetically active tissue during transition from heterotrophic to autotrophic growth. Expression of the blue and red modules was highest in the embryo root and shoot, with expression of the blue module highest in the embryo leaves whilst the difference between the embryonic leaf and root was less notable in red module. Both modules were enriched for chromatin and nucleosome assembly, with the blue module is also enriched for ribosome biogenesis. This is consistent with the known presence of rapidly dividing cells in these meristematic regions. In summary, this analysis was able to correlate functional relationships of genes and their spatial distribution on a sub-tissue level. The grey module was highly expressed in the shoot apical meristem and consisted of genes involved in auxin signalling.

**Figure 6.**
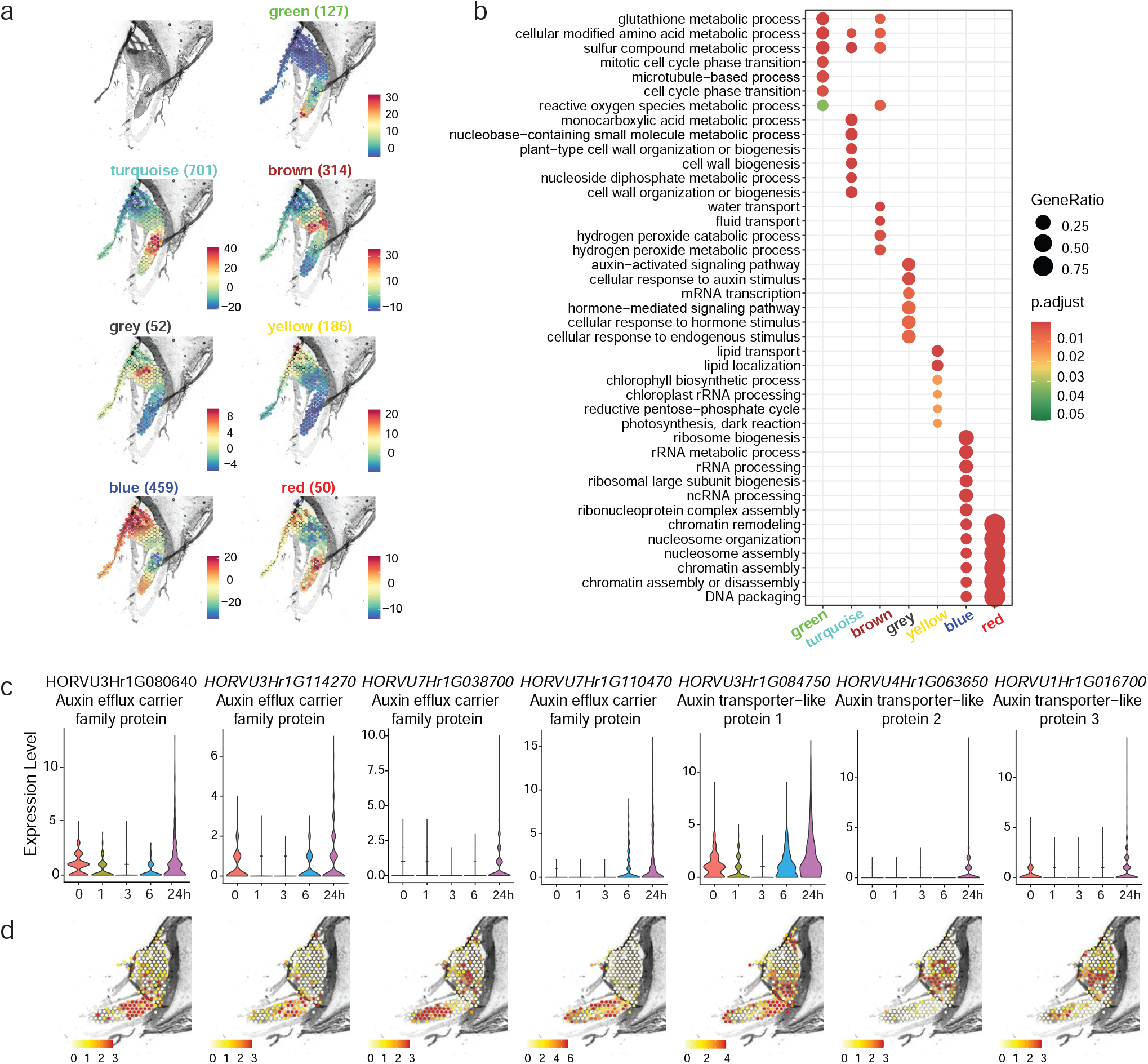
Co-expression of spatially variable genes in the embryo of 24 HAI germinating barley grain. **a**, Module eigengenes, a proxy for expression of genes within each module, of seven co-expressed modules. The top left panel show the bright field microscopic image, the other panels show the overlayed expression data for spots. SVGs showing spatial coherence in embryo of 24h germinating barley grain were grouped into co-expression modules using hdWGCNA^72^. Gene number in each module is given in the brackets. **b**, Gene Ontology enrichment of the co-expression modules. The top six enriched biological process terms for each module are shown. GeneRatio: number of genes annotated with a GO term in a cluster divided by the total number of genes annotated with this GO term. p.adjust: adjusted p value using Benjamini Hochberg method. **c**, Expression of auxin efflux carrier and transport genes showing spatial coherence at 24 HAI. Normalised counts were used to indicate expression level. **d**, Spatial expression of auxin efflux carrier and transport genes showing spatial coherence, colour scales represent normalised counts.

Auxin has a critical role in embryo development and growth^79,80^. An in-depth evaluation of the SVGs list revealed 24 auxin-related genes showing spatially variable expression in at least one time point, with 18 of them found at 24 HAI (Supplementary Table 5). From these 18 auxin-related genes, we further focused on seven genes which belong to auxin efflux carriers and transporter−like families (Fig. 6c). An auxin efflux carrier family protein (*HORVU7Hr1G038700*) and one auxin transporter-like protein (*HORVU4Hr1G063650*) were mainly expressed at 24 HAI while the other five genes were dominantly expressed at more than two time points (Fig. 6c). Interestingly, the four auxin efflux carrier family genes were mainly expressed in the embryo root and enriched towards the basal root (*HORVU3Hr1G080640, HORVU3Hr1G114270*), the inner embryo apex (*HORVU7Hr1G038700*) and epidermis (*HORVU7Hr1G110470*) (Fig. 6d). The three members of auxin transporter−like proteins also display an obvious spatial-dependent expression pattern (Fig. 6d). Auxin transporter−like protein 1 was enriched in the mesocotyl as well as the root tip. Auxin transporter−like protein 2 and 3 are both enriched in the embryo SAM with auxin transporter−like protein 3 also showed high expression in the inner layers of the embryo radicle. The complementary expression of those auxin-related genes suggests the transport and accumulation of auxin in the barley embryo is controlled by a spatially complex distribution of auxin transporters, similar to the well-define role of auxin for the formation of the embryonic apical#x2013;basal axis established in *Arabidopsis*^81^.

## Discussion

Here, we have presented a spatially resolved transcriptomic analysis of germinating barley grain at a resolution of 55 μm over 24 HAI. The accuracy of the spatial determination of expression was confirmed using RNA *in situ* hybridisation. The quantitative nature of the approach shows reasonable correlation with bulk RNA-seq. As different cells will be expressing genes at different levels depending on the cellular landscape^25^, such variation is lost in bulk-sequencing, but retained in spatial transcriptomics. The number of genes detected and that changed in expression in our study was also comparable to bulk sequencing. We developed a web application to allow users to visualize and explore our datasets. The web application gives access to: i) the spatial distribution of clusters in four sections for each time point; ii) expression of genes of interest in a spatial way as well as UMAP and Violin plot; iii) expression of embryo SVGs; iv) marker gene list of each time point and Moran ‘s I value for embryo SVGs. The web application can be accessed at https://spatial.latrobe.edu.au.

In our experiments, we observed consistently isolated individual spots of expression for some genes on single sections. The biological interpretation of such isolated expression is unclear; does it represent transcriptional ‘noise ‘ that has been proposed to occur^82^, or does it represent an artifact? A larger number of replicates will help interpret such signals, and these larger datasets will additionally drive the development of bioinformatic approaches to allow individual cell expression to be visualised and placed in context of the transcriptional landscape of the tissue as a whole. As an advantage for plant biology, the spatial transcriptomic approach works well with fresh frozen tissue. The ability to obtain cell sections from a large variety of plant tissues even with limited amounts means that profiling of cells that have been previously lost either in bulk or protoplast approaches because of their limited numbers will be avoided. A cost limitation currently of this technology is to develop a 3D cell expression atlas of the target tissue. This would require at least tens of sections at each time point. With different commercial technologies emerging in this area and uptake cost should come down in the future^83^, but large scientific consortia like the Plant Cell Atlas (https://www.plantcellatlas.org) and STOC (Spatio-Temporal Omics Consortium -https://www.sto-consortium.org/news_events.html) will be valuable in providing access to individuals to larger funded studies.

From our data the spatial heterogeneity in expression of gene family members and their gradients of expression over time were apparent. The analyses revealed that significant enrichment of different processes occurs in specific locations, e.g. translation in the embryo (Fig. 3), auxin signalling pathways in the coleorhiza (Fig. 6). It is well established that translation, not transcription, is essential for germination to commence. This study shows that this is confined to the embryo, and that 45 genes encoding proteins associated with translation are embryo specific. A variety of genes in other tissues that are associated with grain or processing quality, gamma gliadin, can also now be located to specific areas and promoters associated with this expression defined. Auxin has emerged as a key player in regular pattern for the formation of lateral roots with oscillations in auxin defining the lateral root pattern^84-86^. An auxin maximum precedes lateral root formation which depends on auxin transport. Furthermore, the repression of lateral roots in response to loss of contact with water is regulated by movement of ABA from the phloem^87^. Here, we show that genes encoding auxin transporters already show spatially distinct patterns at the time the radicle emerges from the seed coat. Knowing what auxin transporter(s) (and other hormone transporters) are expressed where and how they respond to environmental changes will allow breeding strategies to target these processes more efficiently. Previously it has been shown that auxin represses aquaporin expression to allow lateral root emergence. The spatial and temporal expression patterns determined here can identify the molecular components involved in this process^88^ and provide approaches to alter these responses for agricultural purposes.

Comparison of marker genes defined using spatial transcriptomics in this study to orthologs in *Arabidopsis* revealed that while some *Arabidopsis* genes had similar a same expression pattern on a tissue basis, the majority did not. This is similar to studies using scRNA-seq showing he expression of some genes in meristems (root or shoot apical meristem) were conserved, and, in general, patterns are conserved across species when similar functions and structures are required^9^. Approximately 65% of genes in plant genomes are duplicates and most crops are polyploid, consequently for many important agronomic traits such as seed size and disease resistance the spatial expression pattern of genes arising from subfunctionalisation and subneofunctionalisation is an important mechanism for retaining duplicates^89^. Thus, determination of the precise sub-tissue expression pattern of genes is key to using molecular approaches for plant improvement in the future.

It is essential that crop varieties that respond to environmental changes are developed to produce the food for a growing world population. While the cessation of growth under environmental limitations has been traditionally viewed as a resource allocation problem, e.g. resources are used for pathogen defence not growth, detailed molecular studies have revealed that growth limitation is due to antagonistic transcriptional pathways^90^. For example, mutation in *PHYTOCHROME B* rescues the growth retardation of jasmonate mutants that have a constitutively active defence response, but this depends on the levels of the defence response and tryptophan biosynthesis^91^. It has also been shown that different stresses can be mediated via different tissue types^92^. Thus, the understanding of where genes are expressed and how they respond to environmental conditions is vital to future strategies to breed plants that can respond to the environment and maintain growth^93^. The concept of specialised plastids is proposed for sensing and responding to stress in a tissue specific manner^94^. Spatial transcriptomics provides the high throughput technique to allow these emerging concepts to be developed and further our knowledge of how plants respond to the environment from a cellular to whole plant level.

## Methods

### Spatial transcriptomics tissue preparation

#### Plant growth and tissue preparation

Barley grains (cv La Trobe) were surface sterilized using the chlorine-gas method in a fume hood, germinated on top of two layers of sterilized filter paper with 12 ml of sterile water added and wrapped with foil at 23 °C. Seeds were harvested at 0, 1, 3, 6 and 24 HAI.

#### Grain sectioning

The seeds for the Visium experiment were collected at the indicated time-points and hand-dissected longitudinally. The seeds were immediately snap frozen in an isopentane bath (2-methylbutane, Sigma Aldrich, cat no. 270342-1L) for 1 min, subsequently embedded in Optimal Cutting Temperature (OCT, Tissue-Tek, cat no. 4583) and the blocks frozen in an isopentane bath and kept on dry ice. The cryoblocks were cut on a cryostat (Leica Biosystems; MA, USA) to a thickness of eight µm at -18°C. The sections were carefully place on a prechilled Tissue Optimization (TO) or Gene Expression (GE) slide and kept frozen at - 80 °C until required.

#### Fixation, staining, and imaging

The slides were dried on a metal plate (PN-1000317, 10x Genomics) in the thermocycler at 37 °C for 1 min, fixed with chilled methanol at -20°C for 30 min, stained with 0.1% (w/v) Safranin O (Sigma-Aldrich, cat no. S8884-25G) in 50% (vol/vol) Ethanol with 2 U/μl RNase inhibitor at room temperature (RT) for 5 min and washed in increasing concentrations of EtOH until the discarded liquid was clear. After drying at 37°C for 1 min, the slides were mounted in 85% (v/v) glycerol + 2 U/μl RNase inhibitor, covered with a coverslip and images of sections were taken using Axio-Imager.M2 microscope (Zeiss). TO slides were captured in on image while GE slide capture areas were imaged individually. Raw images were stitched together using Zen blue software v2.5 (Zeiss). Prior to imaging, the microscope settings were validated using the Visium Imaging Test Slide (PN-2000235). The optimised settings were saved into the TO macro for following experiments. After imaging, the coverslip was removed immediately, and the slide immersed at 45° angle in 3x SSC buffer and washed once with 3x SSC buffer before air drying.

#### Tissue pre-permeabilization, permeabilization and reverse transcription

To pre-permeabilize the tissue, the slides were mounted in a plastic cassette and sections incubated in pre-permeabilization solution^21^ (48 μl Exonuclease I buffer, NEB, cat no. B0293S; 4.5 µl of BSA, Sigma-Aldrich, cat no. A7039-100G; and 2% (w/v) PVP40, Sigma-Aldrich, cat no. PVP40-1KG) at 37 °C for 30 min. This was followed with a wash with 0.1 × SSC buffer (Sigma-Aldrich, cat no. S6639L). The sections were permeabilized with Permeabilization mix™ (10x Genomics) at 37°C for different times (1, 3, 6, 15, 30 min TO slides) or for 3 min (GE slides). Then, wells were washed with 0.1x SSC buffer. After permeabilization, Reverse transcription mixture™ (10x Genomics) was added to each section and incubated at 56 °C for 45 min as described in the 10x Genomics User guide (PN-1000186, CG000239_VisiumSpatialGeneExpression_UserGuide_RevD).

#### Tissue removal and washes (TO slide only)

To remove the tissue, a hydrolytic enzyme mixture was prepared by adding 70 µl of each enzyme (Supplementary Table 7); cellulase (Yakult -”ONOZUKA” R-10, cat no. YAKL0012), pectate lyase (Megazyme, cat no. E-PCLYAN2), xyloglucanase (Megazyme, cat no. E-XEGP), endo 1,4 β-xylanase (Megazyme, cat no. E-XYNACJ), endo 1,4 β-mannanase (Megazyme, cat no. E-BMACJ) and lichenase (Megazyme, cat no. E-LICHN) to 140 µl of 250 mM sodium citrate buffer (Sigma-Aldrich, cat. no. 71497-250 G), pH 6.6. The enzymatic mixture was added to the wells and incubated in a Thermo Mixer at 37 °C for 90 min with shaking (300 r.p.m.). The wells were washed with 0.1x SSC buffer.

Samples were incubated with 10% (v/v) Triton X-100 solution in a Thermo Mixer at 56 °C for 1 h with shaking (300 r.p.m.), followed by a wash with 0.1x SSC buffer. Next wash consisted in a mixture of RLT buffer (Qiagen ref.79216) with 1% (v/v) β-mercaptoethanol, which was incubated in a Thermo Mixer at 56 °C for 1 h with shaking (300 r.p.m.) and followed by a wash with 0.1x SSC buffer. A final incubation with 70 µl proteinase K mixture (60 µl of proteinase K (Qiagen, cat no. 19131) and 420 µl of PKD buffer (Qiagen, cat no. 1034963) was performed in a Thermo Mixer at 56 °C for 1 h with shaking (300 r.p.m.). Hybridization chamber was detached, and the slide washed in a petri dish with 50 °C pre-warmed wash buffer 1 (2x SSC/0.1% SDS) at 50 °C for 10 min with shaking (300 rpm). The slides were further washed with wash buffer 2 (0.2x SSC) and wash buffer 3 (0.1x SSC) at RT for 1 min with shaking (300 rpm). The slide was spin-dried in a swing-bucket centrifuge.

#### Fluorescence imaging or library preparation, and sequencing

When using the Tissue Optimization workflow, a fluorescent image (mRFP filter, Excitation: 559/85 Emission: 600/90) of the cDNA footprint was taken. The whole slide was recorded using a Plan-Apochromat 10X/0.45 M27 objective and the tiling function of the Axio-Imager.M2 microscope (Zeiss). The final file included each of the eight capture areas, 8 × 8 mm (the fiducial frame and the capture area); with the microscope set to ∼1-2 mm beyond the fiducial frame for optimal image alignment. The capture resolution was 0.454 µm/pixel. Raw images were stitched together using Zen blue software v2.5 (Zeiss). When following the Gene Expression workflow, libraries from the different capture areas were synthesized as described in 10x User guide (PN-1000190, CG000239_VisiumSpatialGeneExpression_UserGuide_RevD). Visium libraries were sequenced on an Illumina NextSeq 550 platform according to the 10X Genomics Visium manufacturer ‘s instructions (NextSeq 500/550 High Output kit v2.5 (150 cycles) 20024907, CG000239_VisiumSpatialGeneExpression_UserGuide_RevD), targeting 100 million reads using dual indexing kit TT set A (PN-1000215, 10x Genomics).

### Visium Spatial Gene Expression Analysis

#### Imaging and data processing

Images of the bright field sections were loaded into Loupe browser software (10x Genomics) and the fiducial frames were manually adjusted. The resulting loupe files were used for downstream analysis.

#### Spatial transcriptomics data processing

Selection of spots and image alignment was performed in Loupe Browser (v. 6.2.0, 10x Genomics) to generate alignment files for each section. The sequencing data, bright field images and alignment files were used as input for the Space Ranger (v. 1.3.1, 10x Genomics) to generate gene-spot matrices. IBSCv2 barley genome assembly was used as the reference genome for Space Ranger^95^. For each section, spots with a total UMI count less than 30 and less than 10 expressed genes were excluded from the following analysis. For each time point, the data from four sections were merged with the ‘ ‘merge ‘ ‘ function from Seurat package (v. 4.1.0) and normalized using SCTransform (settings: variable.features.n = NULL, variable.features.rv.th = 1)^12^. Section variability within the data were regressed out using the “vars.to.regress” function from SCTransform. Principal component analysis (PCA) was performed on the genes with residual variance greater than 1, and the 30 most significant components were retained as input for Uniform Manifold Approximation and Projection (UMAP)^96^ and for spot clustering. Generation of spot clusters was made using the “FindNeighbors” (settings: reduction = “pca”, dims = 1:30) followed by the “FindClusters” functions in Seurat, where the default algorithm was used to construct a Shared Nearest Neighbor (SSN) graph and apply the Louvain algorithm for cluster generation. Cluster resolution was set from 0.4 to 0.8. The identity of the clusters was assigned based on their location in the section according to the morphology of a barley grain. To identify marker genes for each cluster, pair-wise comparisons of individual clusters against all other clusters were performed using the “FindAllMarkers” function (settings: min.pct = 0.05, logfc.threshold = 0.25, only.pos = T) with Wilcoxon rank sum test in the Seurat package. The marker genes were further filtered with an adjusted p value < 0.05^22^.

#### Orthology analysis

*Arabidopsis* orthologues to barley genes were identified using the genome pair orthology tool (OMA browser - https://omabrowser.org/oma/genomePW/)^32^. Note that many of the genes were many to many orthologues, but only those within the set that were orthologous to the genes in our subset are shown.

#### Temporal and spatial specific analyses

Supplementary Table 2 show the marker genes of each cluster identified at 0, 1, 3, 6 and 24 HAI. These genes were matched to all the genes in the genome and re-organised such that the marker genes in each of the time points in each of the tissues can be compared across rows (Supplementary Table 4). This revealed that 1,412 genes were identified as marker genes in at least one tissue in at least one time point. In order to unveil any biological processes that may be specific to a tissue, the gene descriptions for each time point for each tissue (25 input sets) were independently examined using a wordcloud generator (https://monkeylearn.com/word-cloud/) to identify the terms occurring multiple times. Note, the words “protein”, “superfamily” and “family” were first removed to avoid bias. Outputs were returned as tables containing terms and number of occurrences for that term in each set. The outputs were then filtered keeping only the terms that were present in the gene-sets from at least three time points for a given tissue or contained five or more genes with that term i.e. spatial expression. Similarly, the terms that were present in the gene-sets from the same time point across at least three tissues were kept i.e. temporal specific expression. In this way, 26 functional descriptions/categories were identified, and these are shown in Fig. 3a.

#### Identification of spatially variable genes and co-expression gene modules

Spots that annotated as embryo tissues were identified for each time point and their expression data normalized using SCTransform as described above^96^. For each section, spatial information between the spots were represented by an adjacency weight matrix which was calculated using the spot coordinates and Moran ‘s I spatial autocorrelation statistics were computed using the normalized expression data along with associated spatial adjacency weight matrix using the MERINGUE framework^97^. SVGs were defined as Moran ‘s I > 0.25, adjusted p value < 0.01, expressed in more than 5% spots in a section, and at least two sections for a time point were considered as for that time point. hdWGCNA was used to perform co-expression analysis for 24 HAI SVGs^72^ using a signed network and ‘bicor ‘ correlation. Module detection was performed with default settings, and the minimal module size was set to 50 genes. After module detection, highly correlated modules were merged using a tree cut height of 0.45. For each module, an eigengene, which is defined as the first principal component of the module ‘s expression pattern, was calculated. The intra-modular connectivity (kME) was calculated using the SignedKME algorithm for each gene, which represents its correlation with the module eigengene value. Gene ontology (GO) annotation of barley were retrieved from Shiny GO database (Version 0.75)^98^ and GO enrichment analysis was performed with the clusterProfiler package^99^. Overrepresentation of GO terms in the category “biological process” was identified by hypergeometric distribution.

### Spatial transcriptomics validation

#### RNA *in situ* hybridization

Seeds were cut longitudinally to equal half and fixed in ice-cold farmers fixative (3:1 ethanol:acetic acid). Samples were placed in the cold room (4 ⁰C) overnight. The fixed tissue was dehydrated using the Leica Semi-Enclosed Benchtop Tissue Processor TP1020 (Leica Biosystems, Mount Waverley, Australia) at room temperature in a graded series of ethanol (1 h each 75%, 85%, 100%, 100%, and 100% v/v). The tissues were then shifted to a graduated ethanol:xylene series (80 min in 75%:25%, 50%:50%, 25%:75% v/v), finished with a xylene series (100% (v/v) twice for 1 h). Tissue was then added to molten Surgipath Paraplast#x00AE; Paraffin (Leica Biosystems) at 65 ⁰C for 2h. Paraplast blocks were then prepared with half seed in each block using the Leica Heated Paraffin Embedding Module EG1150 H with the added Leica Cold Plate for Modular Tissue Embedding System EG1150 C (Leica Biosystems). with vacuum infiltration. Embedded tissues were cut at 8 µm sections and *in situ* hybridization was carried out according to modified protocols from^100^ 50 ⁰C hybridization temperature and 0.2x SSC washes^100^. Gene of interest was amplified using designed primers (Supplementary Table 8) and cloned into pGEMT-Easy vector (Promega). Using the DIG RNA Labelling Kit (Roche Diagnostics), Digoxigenin-labelled antisense and sense RNA probes were transcribed from T7 or SP6 promoter of pGEMT-Easy vector (Promega) according to manufacturer ‘s instructions. All hybridization results were observed and photographed using a Zeiss Axio Observer A1 microscope (Carl Zeiss AG).

### Data Availability

The accession number for the raw sequencing data and Space Ranger processed data reported in this paper is Gene Expression Omnibus (GEO): GSE218970 (https://www.ncbi.nlm.nih.gov/geo/). Any additional information required to reanalyse the data reported in this paper is available upon request.

The interactive web browser can be explored at https://spatial.latrobe.edu.au/ with the username: reviewer and the password: barley_pass.

## Acknowledgements

We thank Dr. Matthew Tucker for kindly providing the barley grains for this experiment. We thank Dr. Jacqueline Orian and her laboratory for their guidance setting up the cryosectioning. Work in M.G.L. ‘s lab is funded by the Australian Research Council (ARC) Discovery Program grant DP220102840. J.W. and L.C.L. were funded by an ARC Discovery Program grant DP210103258. J.W. is supported by a Kun Peng Fellowship from the Zhejiang Provincial government.

## Figure Legends

**Supplementary Figure 1**. Overview of Spatial Transcriptomics experiment on barley germinating grains.

a, Representative graph of barley germination showing images of the barley grains collected at the given time points of the experiment and the main processes occurring during the span of 48 h. Scale bar corresponds to 1 mm. b, Representative Visium capture areas containing a barley grain section at the given time points. Sections correspond to one out of four with 8-µm longitudinal thickness and the embryo in the bottom left corner and the endosperm on the top right. Cryosections at the same time point were collected from the same seed in serial sections. c, Each gene expression (GE) slide contains four capture areas (6.5 × 6.5 mm) defined by an outside fiduciary frame (total area size 8 × 8 mm). Each capture area contains exactly 4992 gene expression spots with spatial primers. These include Illumina TruSeq Read 1 for sequencing, 16 nucleotides (nt) spatial barcode that identifies the spot location within the array, 12 nt unique molecular identifier (UMI) that identifies each mRNA and 30 nt poly(dT) sequence that captures the polyA tail of the mRNA for cDNA synthesis. The spots are 55 µm in diameter and 100 µm centre to centre. d, On-slide Optimized Visium Gene Expression solution workflow: (i) adequate sections of the tissue are prepared using a cryostat, (ii) cryosections are positioned on top of an array covered with primers, (iii) samples are fixed in methanol and stained with Safranin O, (iv) sections are imaged using an Axio-Imager M.2 microscope (Zeiss). Then (v) sections are permeabilized using the permeabilization solution and finally, (vi) cDNA is synthesized to amplify the mRNAs, UMIs and spatial barcodes. Scale bars correspond to 5 mm. e, Off-slide Visium Gene Expression solution workflow consists of library preparation, sequencing and visualization of the data. 10x Genomics Visium solution data are analysed using Space Ranger software and the outputs are visualized using Loupe browser. Scale bars correspond to 5 mm.

**Supplementary Figure 2**. Spatial transcriptomics tissue optimisation protocol for barley grains.

a, For optimization experiments, tissue optimization (TO) slides were used. Each capture area on the slide, eight in total, is defined by an edge. Each area is uniformly coated with oligonucleotides for mRNA capture, the probes have poly(dT) primers to enable cDNA synthesis from the poly-adenylated mRNAs. There are no spatial barcodes in these primers. The TO slides are designed to test the permeabilization conditions for the experiment and it is recommended to use one slide per tissue type. b, The optimized permeabilisation time was defined by visualizing the fluorescent footprint of the cDNA on top of the array. BF corresponds to the brightfield imaging of the fixed, safranin-stained TO slide, RFP, to the fluorescent cDNA footprint after RT reaction and Z, to the white square area in RFP picture. E, starchy endosperm; S, scutellum; E, embryo. Scale bars correspond to 1 mm. c, Visium spatial tissue optimisation workflow. Optimised protocol from 10x Genomics for mammalian tissues. The protocol can be performed in one day. d, Visium Spatial PLANT tissue optimization workflow. First difference between the protocols was the dye used for barley grain staining. Safranin O was determined to be performing better to protect the grain architecture. Tissue removal steps were also modified in this protocol. Tissue removal #1 consisted of a hydrolytic mixture of enzymes (Supplementary Table 7). Tissue removal #2 consisted of several incubation steps. e, Intersect analysis of 24 HAI samples using the two different permeabilization methods: permeabilization (perm) and pre-permeabilization (pre-perm), which shows the number of genes identified by either of the methods are similar.

**Supplementary Figure 3**. UMI number and gene number in different sections.

a, Log transformed UMI number in different sections of 0 HAI (i), 1 HAI (ii), 3 HAI (iii), 6 HAI (iv) and 24 HAI (v). Each dot is a spot. b, Number of genes with UMI > 0 in different sections of 0 HAI (i), 1 HAI (ii), 3 HAI (iii), 6 HAI (iv) and 24 HAI (v). Each dot is a spot.

**Supplementary Figure 4**. UMI number in spatial feature plot across different time points. UMI number of 0 HAI (a), 1 HAI (b), 3 HAI (c), 6 HAI (d) and 24 HAI (e).

**Supplementary Figure 5**. Gene number in spatial feature plot across different time points. Gene number of 0 HAI (a), 1 HAI (b), 3 HAI (c), 6 HAI (d) and 24 HAI (e).

**Supplementary Figure 6**. Spatial transcriptome of 0 HAI germinating barley grain.

a, Spatial visualization of the unbiased clustering of spots for four 0 HAI barley sections. Top left panel, BF microscopic image of section C, top middle panel, spatial localization of each cluster of section C, right panel, merged BF image and spatial clusters of section C. Bottom panels, merged BF image and spatial clusters of other three sections The tissue/cell-type identity of each cluster was assigned based on the location of each cluster. b, Uniform manifold approximation and projection (UMAP) of spatial spots from four 0 HAI barley sections. Dots, individual spots from the Visium slide; n = 4744 spots; colours indicate the cluster number that each spot belongs to. c, UMAP of spatial spots from four 0 HAI barley sections, colours indicate different sections for each spot. d, Expression of the top five marker genes of coleorhiza, embryo, scutellum, aleurone and endosperm. Circle size indicates the percentage of spots expressing the marker and colour represents z-scored expression value. Col: coleorhiza, emb: embryo, scu: scutellum, ale: aleurone, end: endosperm.

**Supplementary Figure 7**. Spatial transcriptome of 1 HAI germinating barley grain.

a, Spatial visualization of the unbiased clustering of spots for four 1 HAI barley sections. Top left panel, BF microscopic image of section D, top middle panel, spatial localization of each cluster of section D, top right panel, merged BF image and spatial clusters of section D. Bottom panels, merged BF image and spatial clusters of other three sections. The tissue/cell-type identity of each cluster was assigned based on the location of each. b, Uniform manifold approximation and projection (UMAP) of spatial spots from four 1 HAI barley sections. Dots, individual spots from the Visium slide; n = 5712 spots; colours indicate the cluster that each spot belongs to. c, UMAP of spatial spots from four 1 HAI barley sections, colours indicate different sections for each spot. **d**, Expression of the top five marker genes of coleorhiza, embryo, scutellum, aleurone and endosperm. Circle size indicates the percentage of spots expressing the marker and colour represents z-scored expression value. Col: coleorhiza, emb: embryo, scu: scutellum, ale: aleurone, end: endosperm.

**Supplementary Figure 8**. Spatial transcriptome of 3 HAI germinating barley grain.

a, Spatial visualization of the unbiased clustering of spots for four 3 HAI barley sections. Top left panel, BF microscopic image of section A, top middle panel, spatial localization of each cluster of section A, top right panel, merged BF image and spatial clusters of section A. Bottom panels, merged BF image and spatial clusters of other three sections. The tissue/cell-type identity of each cluster was assigned based on the location of each cluster. b, Uniform manifold approximation and projection (UMAP) of spatial spots from four 3 HAI barley sections. Dots, individual spots from the Visium slide; n = 5881 spots; colours indicate the cluster number that each spot belongs to. c, UMAP of spatial spots from four 3 HAI barley sections, colours indicate different sections for each spot. d, Expression of the top five marker genes of coleorhiza (col), embryo (emb), scutellum (scu), aleurone (ale) and endosperm (end). Circle size indicates the percentage of spots expressing the marker and colour represents z-scored expression value. Col: coleorhiza, emb: embryo, scu: scutellum, ale: aleurone, end: endosperm.

**Supplementary Figure 9**. Spatial transcriptome of 6 HAI barley grain.

a, Spatial visualization of the unbiased clustering of spots for four 6 HAI barley section. top left panel, BF microscopic image of section A, top middle panel, spatial localization of each cluster of section A, top right panel, merged BF image and spatial clusters of section A. Bottom panels, merged BF image and spatial clusters of other three sections. The tissue/cell-type identity of each cluster was assigned based on the location of each cluster. b, Uniform manifold approximation and projection (UMAP) of spatial spots from four 6 HAI barley sections. Dots, individual spots from the Visium slide; n = 4172 spots; colours indicate the cluster number that each spot belongs to. c, UMAP of spatial spots from four 6 HAI barley sections, colours indicate different sections for each spot. d, Expression of the top five marker genes of coleorhiza, embryo, scutellum, aleurone and endosperm. Circle size indicates the percentage of spots expressing the marker and colour represents z-scored expression value. Col: coleorhiza, emb: embryo, scu: scutellum, ale: aleurone, end: endosperm.

**Supplementary Figure 10**. Barley grain anatomy and tissue annotations at 24 HAI.

a, Tissue architecture of a 24 HAI barley grain cv La Trobe. b, Tissue annotations based on the histology of the 24 HAI barley grain.

**Supplementary Figure 11**. Correlation between different sections at each time point.

Pair-wise Pearson correlation between different sections at 0 HAI (a), 1 HAI (b), 3 HAI (c) and 6 HAI (d). Dots correspond to genes expressed in at least 5 spots. X axis and Y axis show the average of normalized count of tissue covered spots from two different sections, respectively. Pearson correlation coefficient is indicated as *R*.

**Supplementary Figure 12**. Percentage of spots of four barley sections for each cluster for different time point.

Percentage of spots of four barley sections for each cluster for 0 HAI (a), 1 HAI (b), 3 HAI (c), 6 HAI (d) and 24 HAI (e). The number of spots were also shown in the parentheses.

**Supplementary Figure 13**. Cluster correlation across different sections of 0 HAI.

Pair-wise Pearson correlation of different clusters across different sections at 0 HAI. Genes with UMI > 0 in at least 5 spots were used in this analysis. For each gene, the average of normalized count of all spots per cluster and section were used to calculate the Pearson correlation coefficient. A, B, C and D indicate different sections. The same cluster from four sections were shown in a black box.

**Supplementary Figure 14**. Cluster correlation across different sections of 1 HAI.

Pair-wise Pearson correlation of different clusters across different sections at 1 HAI. Genes with UMI > 0 in at least 5 spots were used in this analysis. For each gene, the average of normalized count of all spots per cluster and section were used to calculate the Pearson correlation coefficient. A, B, C and D indicate different sections. The same cluster from four sections were shown in a black box.

**Supplementary Figure 15**. Cluster correlation across different sections of 3 HAI.

Pair-wise Pearson correlation of different clusters across different sections at 3 HAI. Genes with UMI > 0 in at least 5 spots were used in this analysis. For each gene, the average of normalized count of all spots per cluster and section were used to calculate the Pearson correlation coefficient. A, B, C and D indicate different sections. The same cluster from four sections were shown in a black box.

**Supplementary Figure 16**. Cluster correlation across different sections of 6 HAI.

Pair-wise Pearson correlation of different clusters across different sections at 6 HAI. Genes with UMI > 0 in at least 5 spots were used in this analysis. For each gene, the average of normalized count of all spots per cluster and section were used to calculate the Pearson correlation coefficient. A, B, C and D indicate different sections. The same cluster from four sections were shown in a black box.

**Supplementary Figure 17**. Cluster correlation across different sections of 24 HAI.

Pair-wise Pearson correlation of different clusters across different sections at 24HAI. Genes with UMI > 0 in at least 5 spots were used in this analysis. For each gene, the average of normalized count of all spots per cluster and section were used to calculate the Pearson correlation coefficient. A, B, C and D indicate different sections. The same cluster from four sections were shown in a black box.

**Supplementary Figure 18**. Correlation of gene expression in Spatial transcriptomics and tissue-specific RNA-seq experiments.

Gene expression of genes detected across comparable tissues in the Spatial transcriptomics experiment and a tissue-specific RNA-seq experiment^24^ were correlated. Only genes with expression above 5 UMI or 5 TPM, respectively, were included. To allow for the comparison of the two experiments, gene expression was averaged across spots of the same tissue for the Spatial transcriptomics experiment and across the three aleurone sections of the tissue-specific RNA-seq experiment. The coefficient of determination (R²) for the linear regression is indicated in each panel.

**Supplementary Figure 19**. UMI number of different clusters across different time points.

UMI number of different clusters at 0 HAI (a), 1 HAI (b), 3 HAI (c), 6 HAI (d) and 24 HAI (e). Col: coleorhiza, emb: embryo, scu: scutellum, ale: aleurone, end: endosperm.

**Supplementary Figure 20**. Gene number of different clusters across different time points.

Gene number of different clusters at 0 HAI (a), 1 HAI (b), 3 HAI (c), 6 HAI (d) and 24 HAI (e). Col: coleorhiza, emb: embryo, scu: scutellum, ale: aleurone, end: endosperm.

**Supplementary Figure 21**. Analysis of the tissue expression pattern of the *Arabidopsis thaliana* orthologs of the top 10 marker genes per cluster for endosperm at each time point. The Arabidopsis orthologs indicated in red displayed a similar tissue expression pattern (green box) to the barley ortholog.

**Supplementary Figure 22**. Analysis of the tissue expression pattern of the *Arabidopsis thaliana* orthologs of the top 10 marker genes per cluster for aleurone at each time point. The Arabidopsis orthologs indicated in red displayed a similar tissue expression pattern green box) to the barley ortholog.

**Supplementary Figure 23**. Analysis of the tissue expression pattern of the *Arabidopsis thaliana* orthologs of the top 10 marker genes per cluster for scutellum at each time point. The Arabidopsis orthologs indicated in red displayed a similar tissue expression pattern (green box) to the barley ortholog.

**Supplementary Figure 24**. Analysis of the tissue expression pattern of the *Arabidopsis thaliana* orthologs of the top 10 marker genes per cluster for embryo at each time point. The Arabidopsis orthologs indicated in red displayed a similar tissue expression pattern (green box) to the barley ortholog.

**Supplementary Figure 25**. Summary of the orthologous genes from *Arabidopsis* and barley that show similar tissue specific expression.

**Supplementary Figure 26**. Transport-related gene expression during barley germination.

Bubble plot of transport-related genes highly expressed during barley germination. Circle size indicates the percentage of expression and colour indicates average expression after z-scoring. Col: coleorhiza, emb: embryo, scu: scutellum, ale: aleurone, end: endosperm.

**Supplementary Figure 27**. Cell wall-related gene expression during barley germination.

Bubble plot of cell wall-related genes highly and specifically expressed during barley germination. Circle size indicates the percentage of expression and colour indicates average expression after z-scoring. Col: coleorhiza, emb: embryo, scu: scutellum, ale: aleurone, end: endosperm. Coloured HORVU IDs correspond to spatial images in Supplementary Fig. 19.

**Supplementary Figure 28**. Spatio-temporal expression of cell wall protein-related genes during barley germination. a, Spatio-temporal expression of *HORVU3Hr1G073780* gene, a glucan synthase, in charge of cellulose biosynthesis was spatially expressed in the outer layers of the radicle at 24 HAI. b, Spatio-temporal expression of *HORVU3Hr1G081180* gene, an alpha-expansin expressed during later germination and involved in cell wall modification. c, Spatio-temporal expression of *HORVU5Hr1G080860* gene, a glucose epimerase related to carbohydrate precursor synthesis, expressed in scutellum across all time points during germination. Black arrows indicate areas of expression. Colour scales represent normalised counts.

**Supplementary Figure 29**. Transcription factor-related gene expression during barley germination. Bubble plot of transcription factor-related genes highly expressed during barley germination. Circle size indicates the percentage of expression and colour indicates average expression after z-scoring. Col: coleorhiza, emb: embryo, scu: scutellum, ale: aleurone, end: endosperm.

**Supplementary Figure 30**. Spatio-temporal expression of transcription factors during barley germination. a, Bubble plot showing the spatially resolved expression of several transcription factor encoding genes. Circle size indicates the percentage of expression and colour indicates average expression after z-scoring. The x-axis represents individual clusters identified across tissues. **b**, Spatial expression of transcription factor genes, (i) Karrikin-responsive bZIP (ii) *BARLEY KNOX3* (*BKN3)*, (iii) WUSCHEL-related homeobox, (iv) bHLH PHY-related and (v) AP2/EREB RAP2.9. Colour scales represent the normalised counts with the same cut-off in each row or with cut-offs individually specified on the image. Black arrows indicate areas of expression. Col: coleorhiza, emb: embryo, scu: scutellum, ale: aleurone, end: endosperm.

**Supplementary Figure 31**. Overlap of number of SVGs in different sections across different time points.

Number of SVGs in different sections at 0 HAI (a), 1 HAI (b), 3 HAI (c), 6 HAI (d) and 24 HAI (e). Yellow: genes with Moran ‘s I > 0.25, adjusted p value < 0.01, expressed in more than 5% spots in at least two sections, and the total number were boxed with yellow and plotted in Figure 5a. Blue, genes with Moran ‘s I > 0.25, adjusted p value < 0.01, expressed in more than 5% spots in only one section.

## Supplemental Tables

**Supplemental Table 1**. Sequencing data quality metrics.

**Supplemental Table 2**. Cluster marker gene of all datasets.

**Supplemental Table 3**. Top 10 marker genes for all clusters at all time points.

**Supplemental Table 4**. Marker genes of each identified cluster in 0 HAI, 1 HAI, 3 HAI, 6 HAI and 24 HAI germination barley seed.

**Supplemental Table 5**. Spatial heterogeneity of embryo expressed genes.

**Supplemental Table 6**. Co-expression module membership of 24 HAI significant spatially variable genes.

**Supplemental Table 7**. Hydrolytic enzymes used in tissue removal step.

**Supplemental Table 8**. *In situ* hybridisation sequencing primer list.

