## Supplementary material for "Spatially Resolved Transcriptomic Analysis of the Germinating Barley Grain": Supp Fig 1

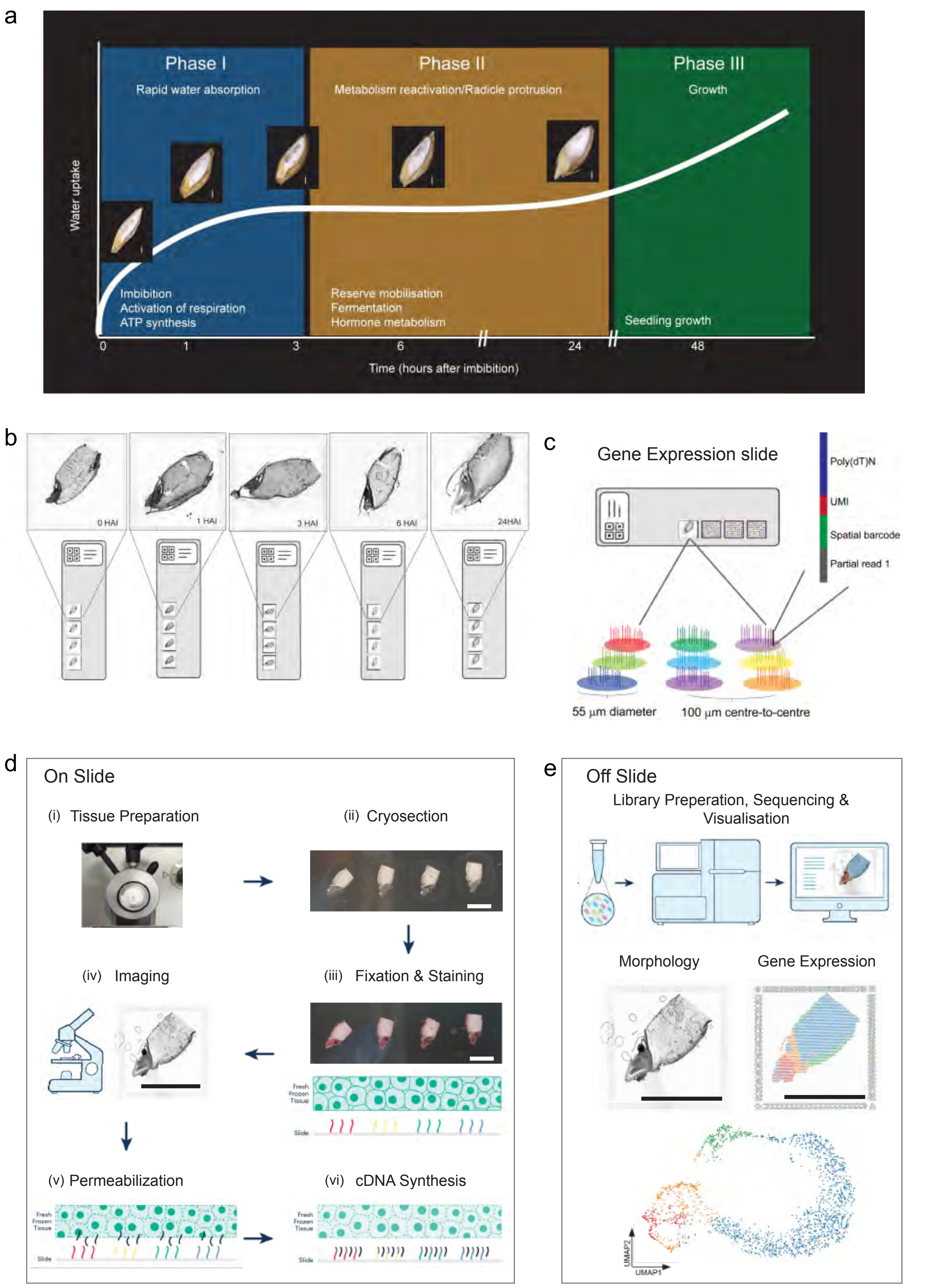


**Supplementary Figure 1.** Overview of Spatial Transcriptomics experiment on barley germinating grains.
