## Supplementary material for "Spatially Resolved Transcriptomic Analysis of the Germinating Barley Grain": Supp Fig 2

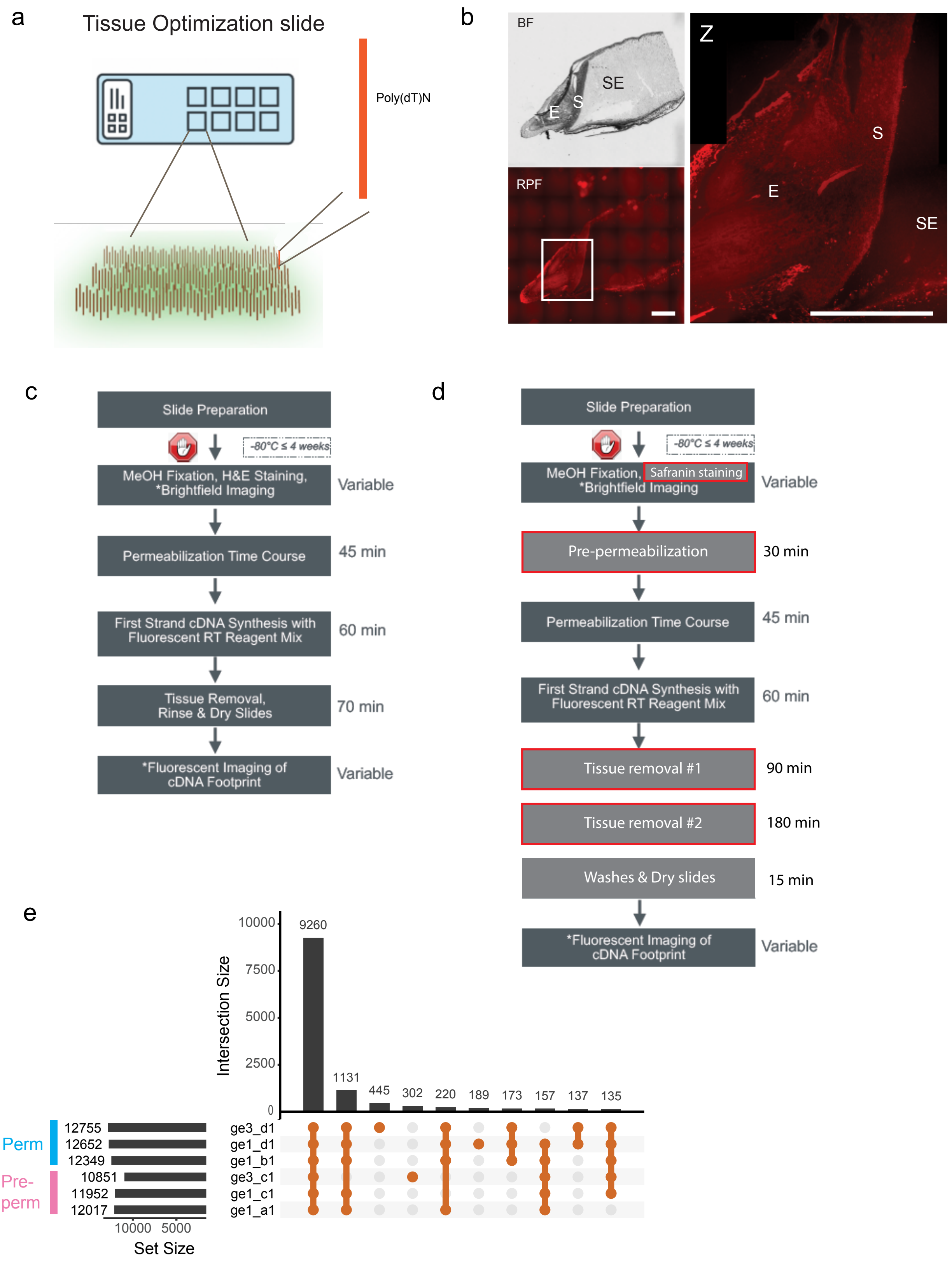


**Supplementary Figure 2.** Spatial transcriptomics tissue optimisation protocol for barley grains.
