## Supplementary figures and images for "Spatially Resolved Transcriptomic Analysis of the Germinating Barley Grain"

### Supp Fig 3

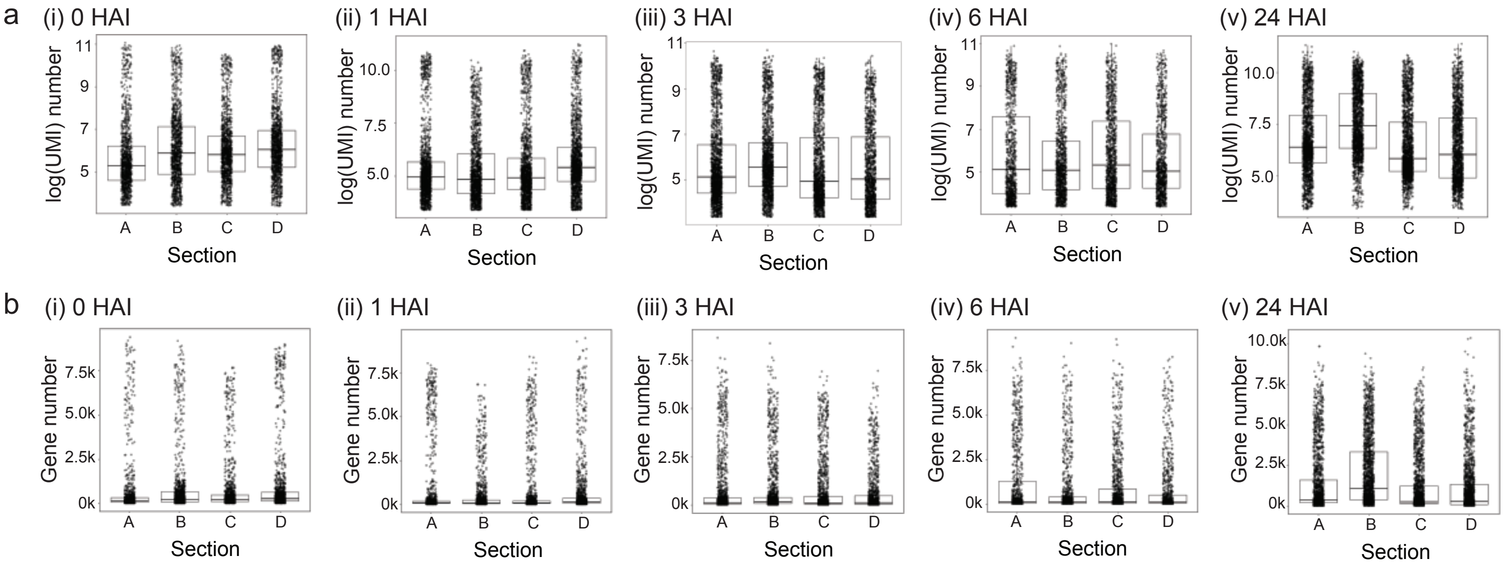


**Supplementary Figure 3.** UMI number and gene number in different sections.

### Supp Fig 4

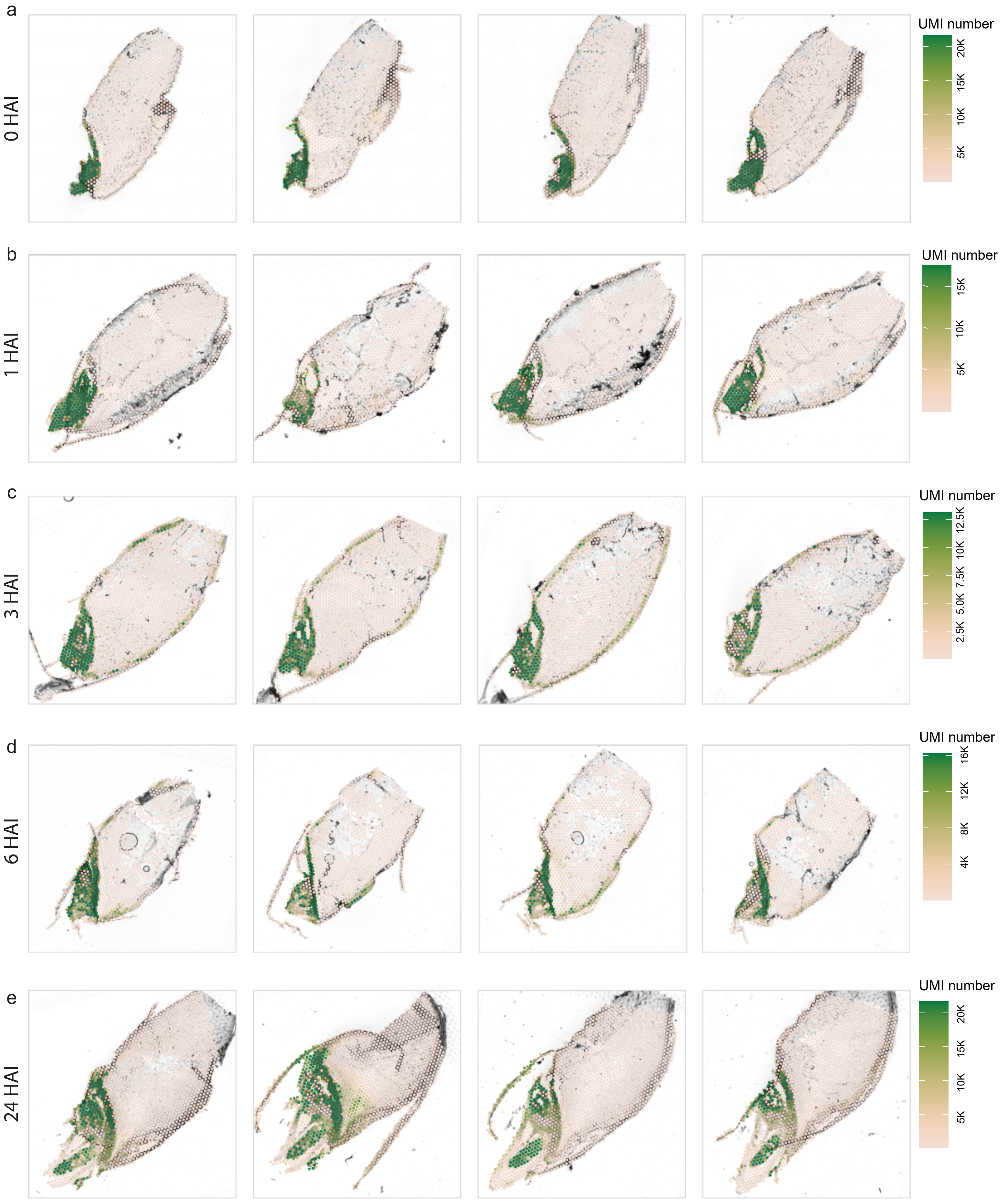


**Supplementary Figure 4.** UMI number in spatial feature plot across different time points.

### Supp Fig 5

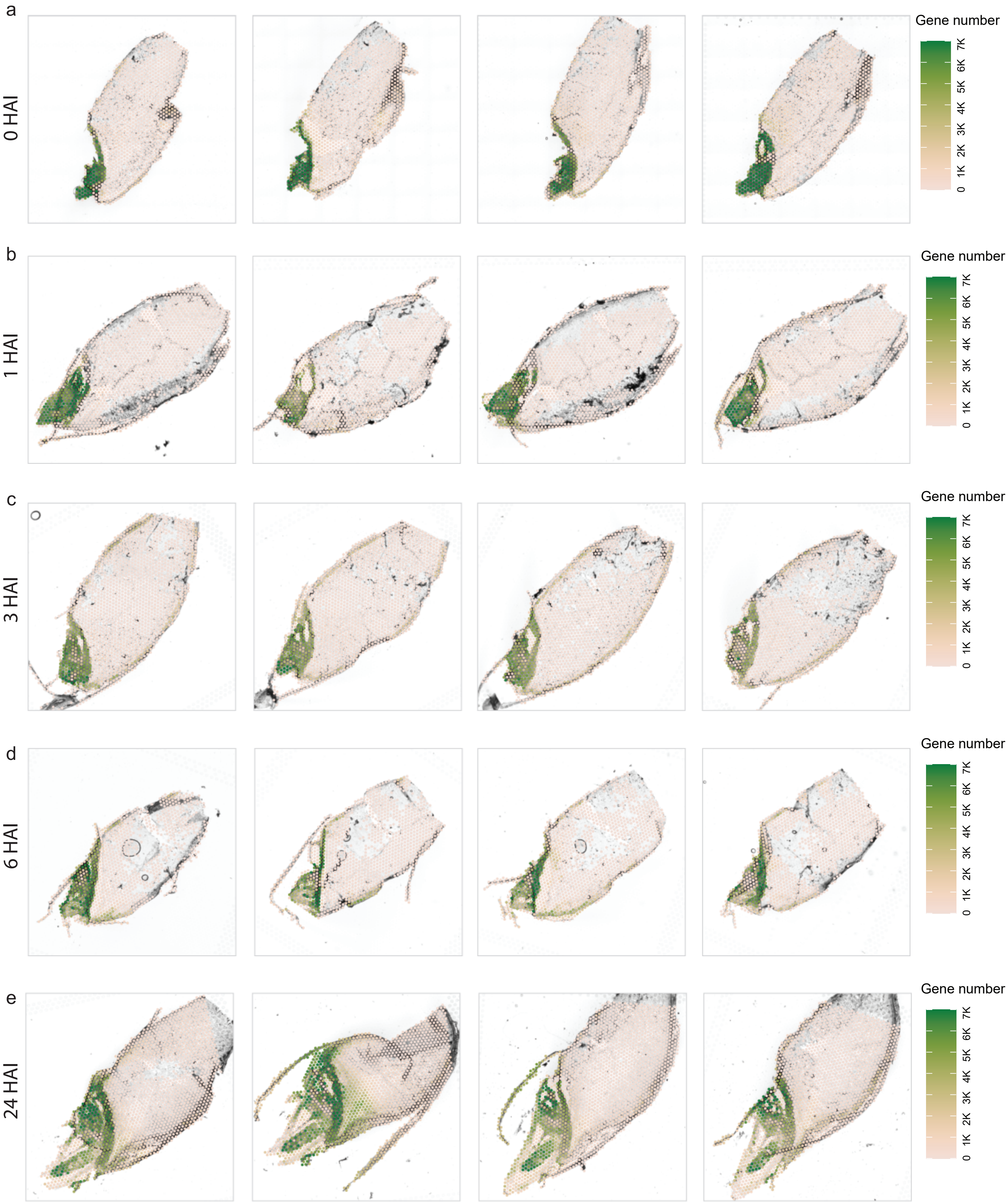


**Supplementary Figure 5.** Gene number in spatial feature plot across different time points.

### Supp Fig 6

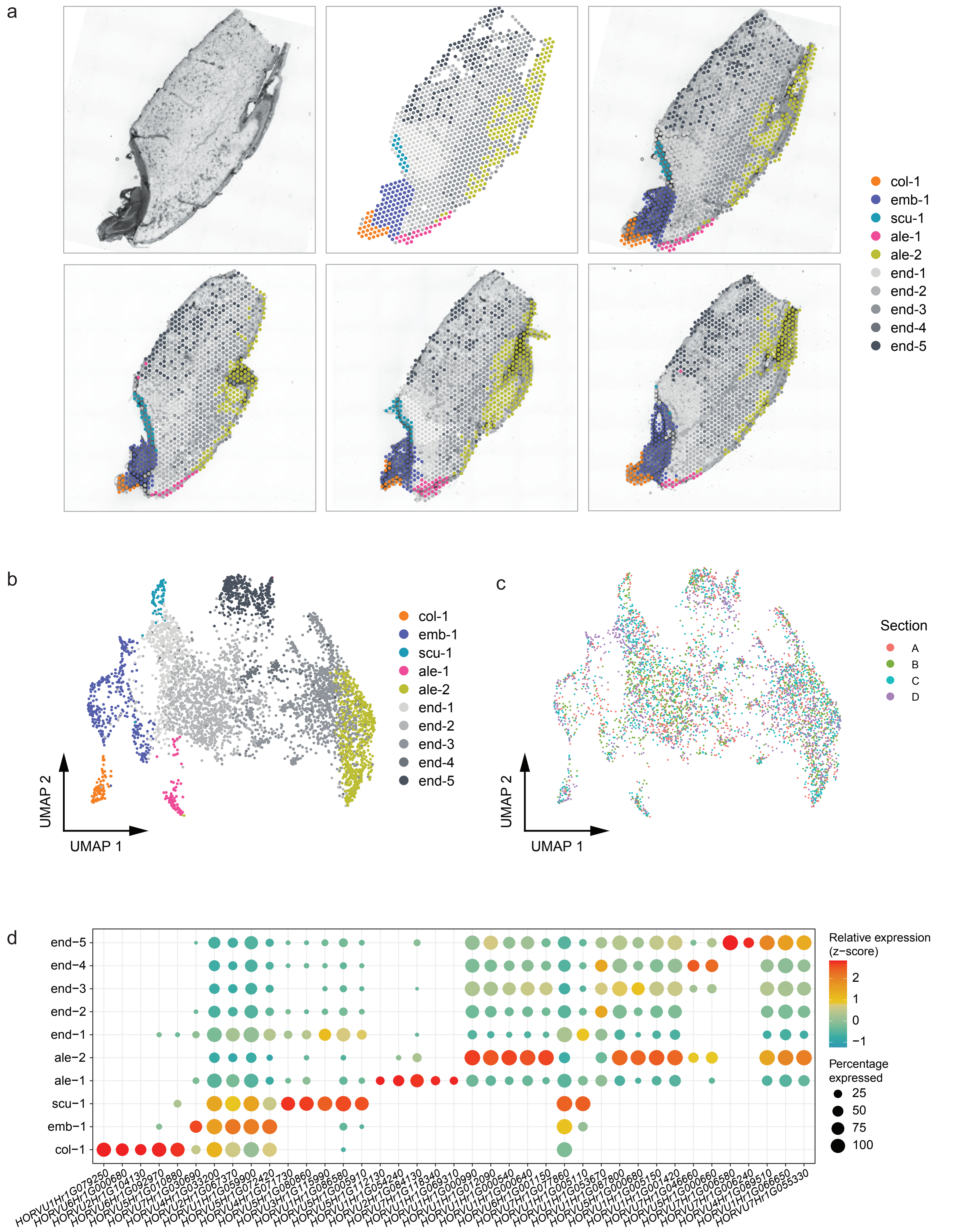


**Supplementary Figure 6.** Spatial transcriptome of 0 HAI germinating barley grain.

### Supp Fig 7

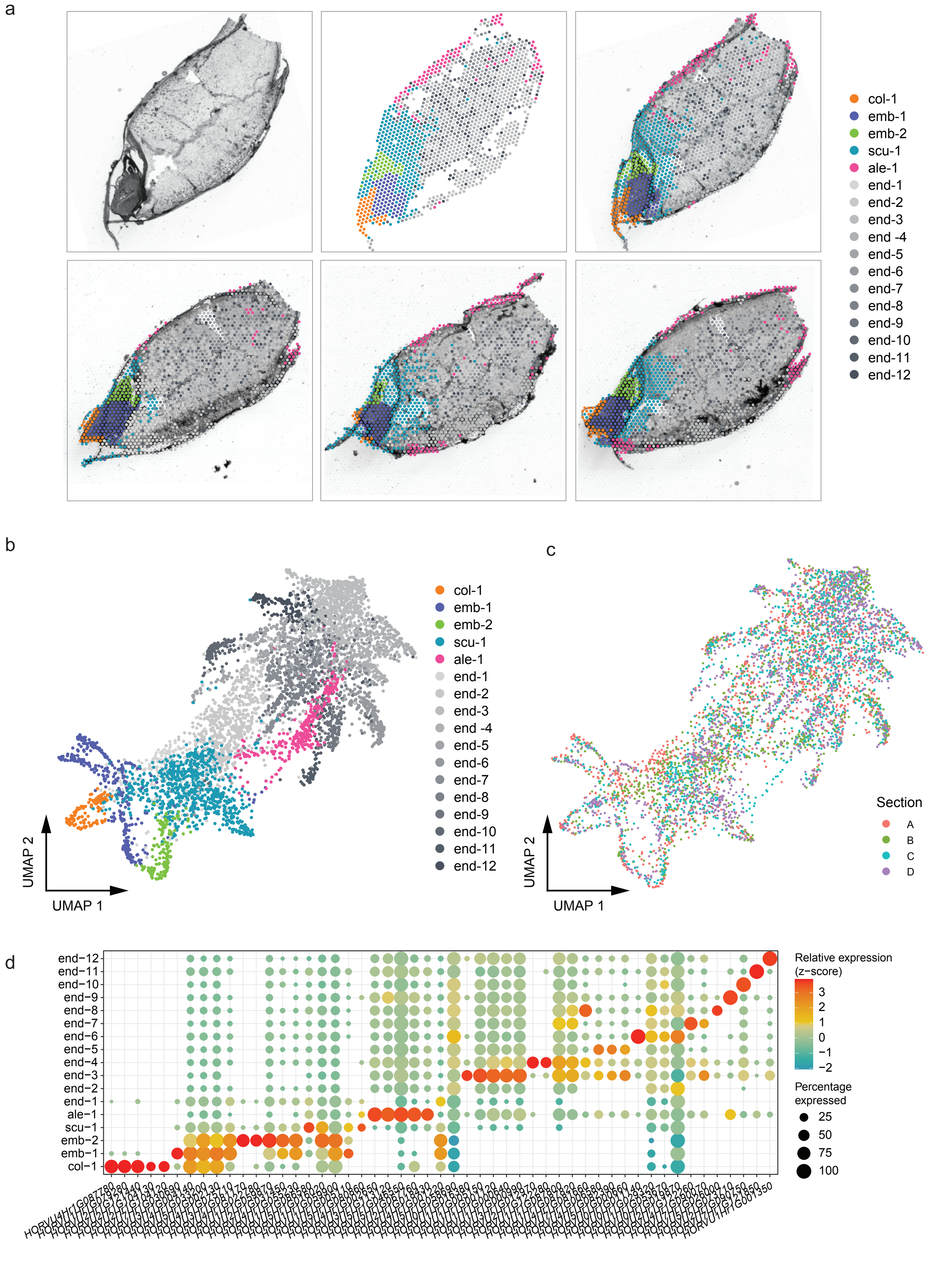


**Supplementary Figure 7.** Spatial transcriptome of 1 HAI germinating barley grain.

### Supp Fig 8

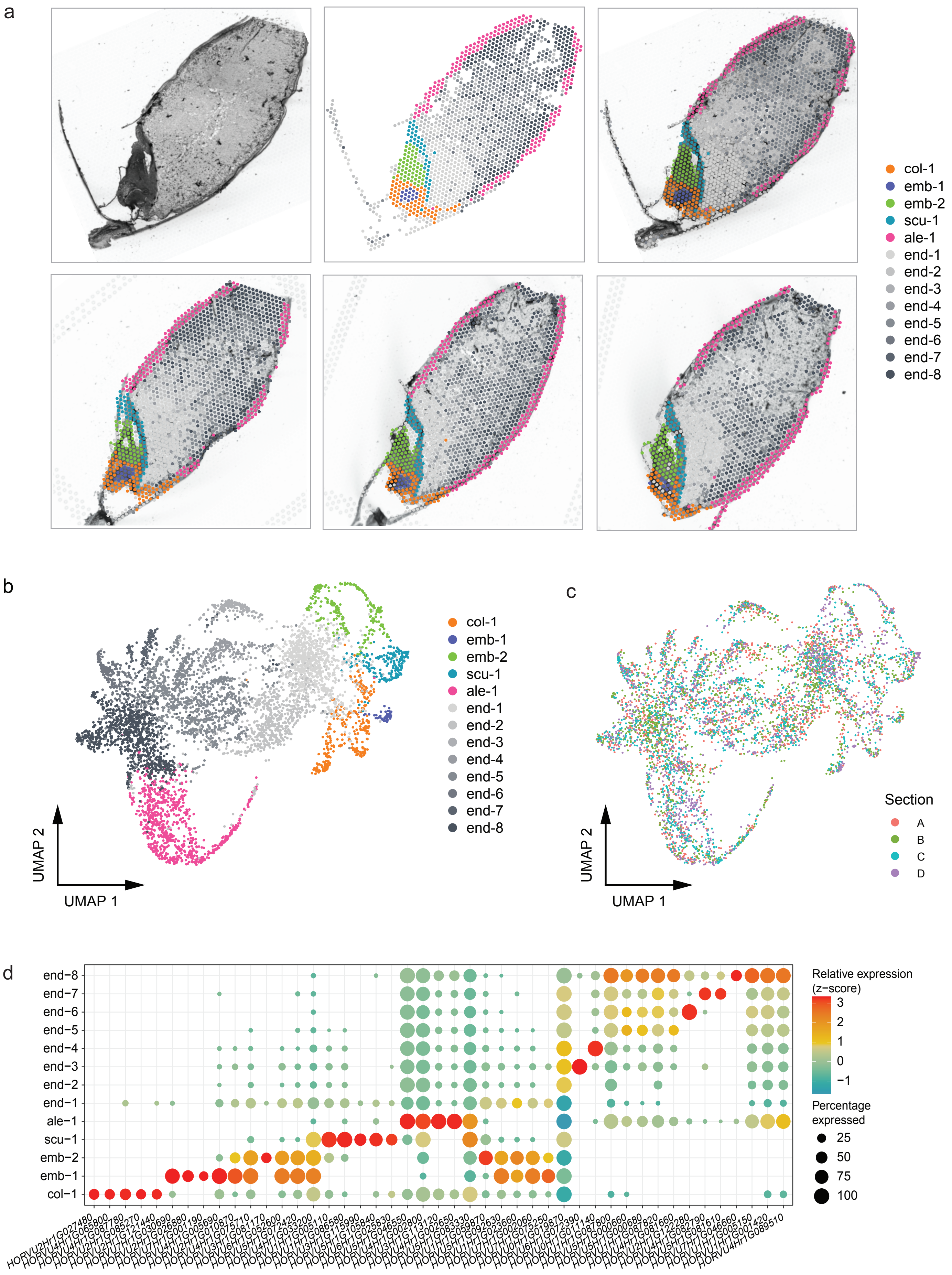


Supplementary Figure 8. Spatial transcriptome of 3 HAI germinating barley grain.

### Supp Fig 9

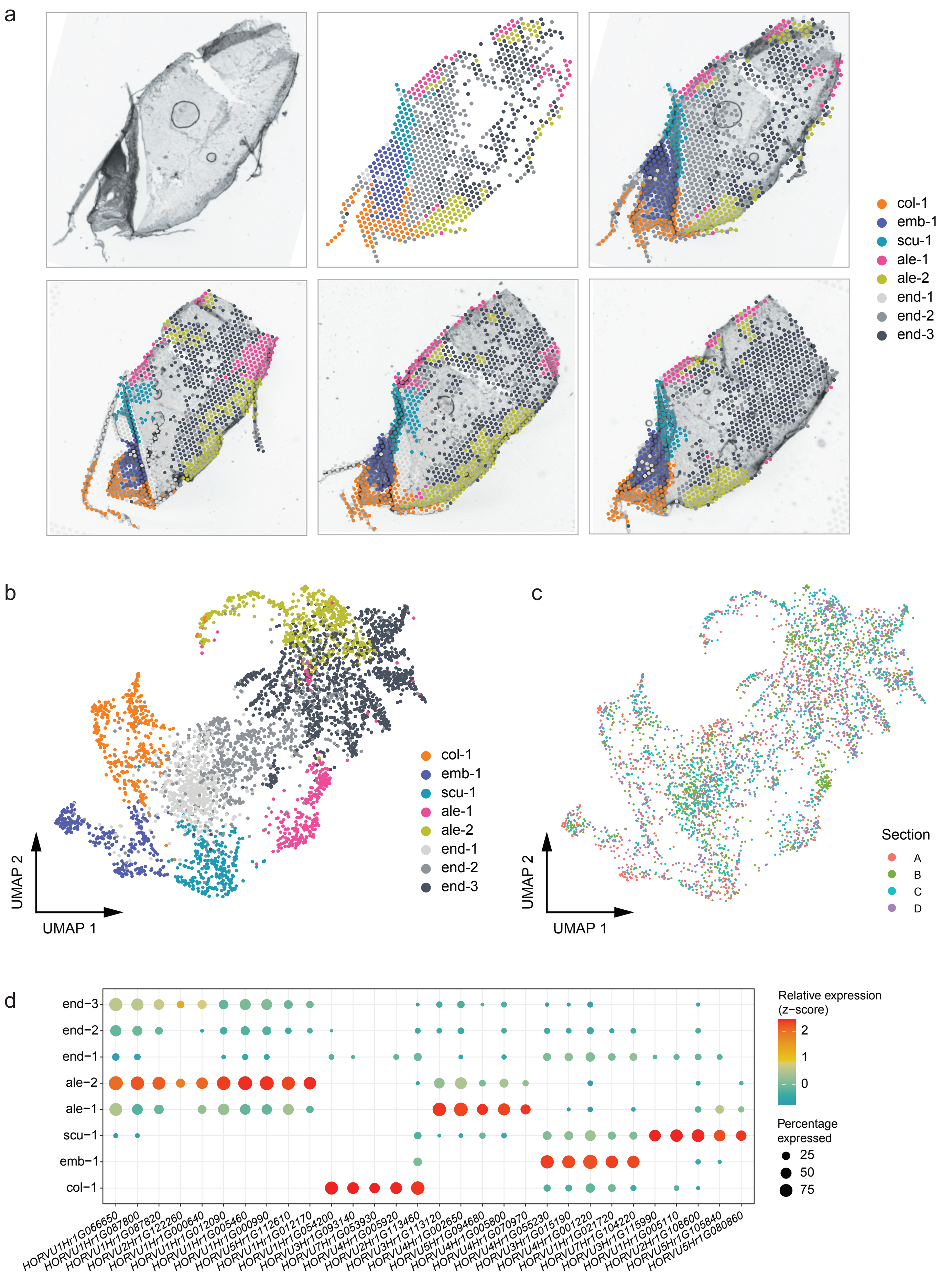


**Supplementary Figure 9.** Spatial transcriptome of 6 HAI barley grain.

### Supp Fig 10

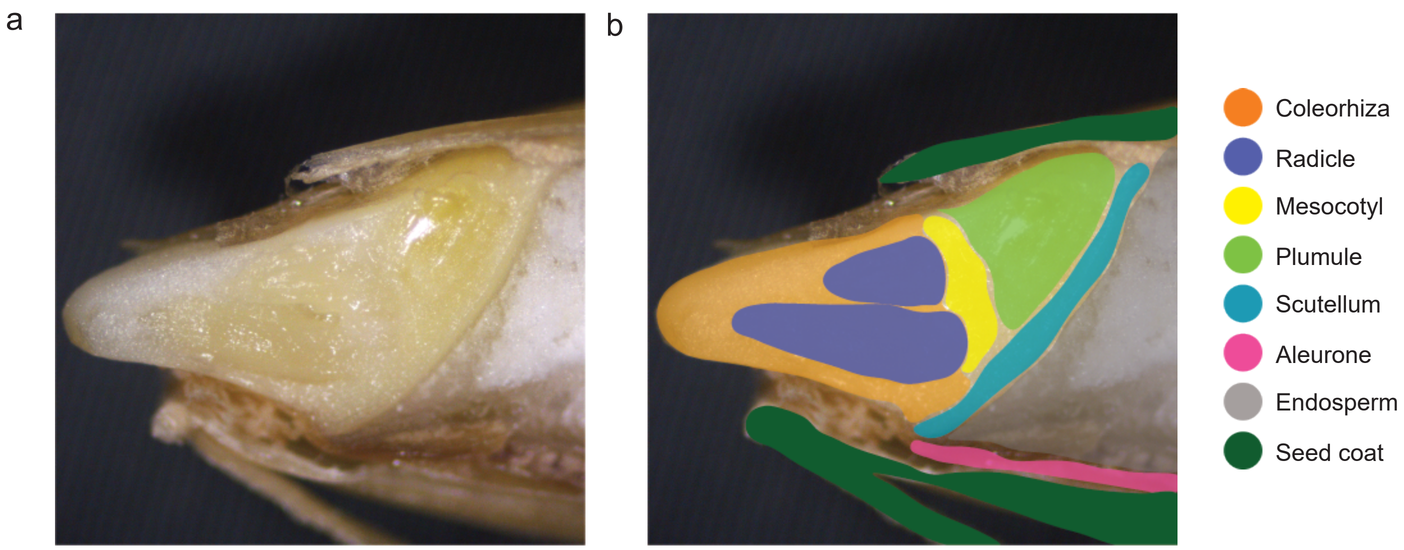


**Supplementary Figure 10.** Barley grain structures and annotations at 24 HAI.

### Supp Fig 11

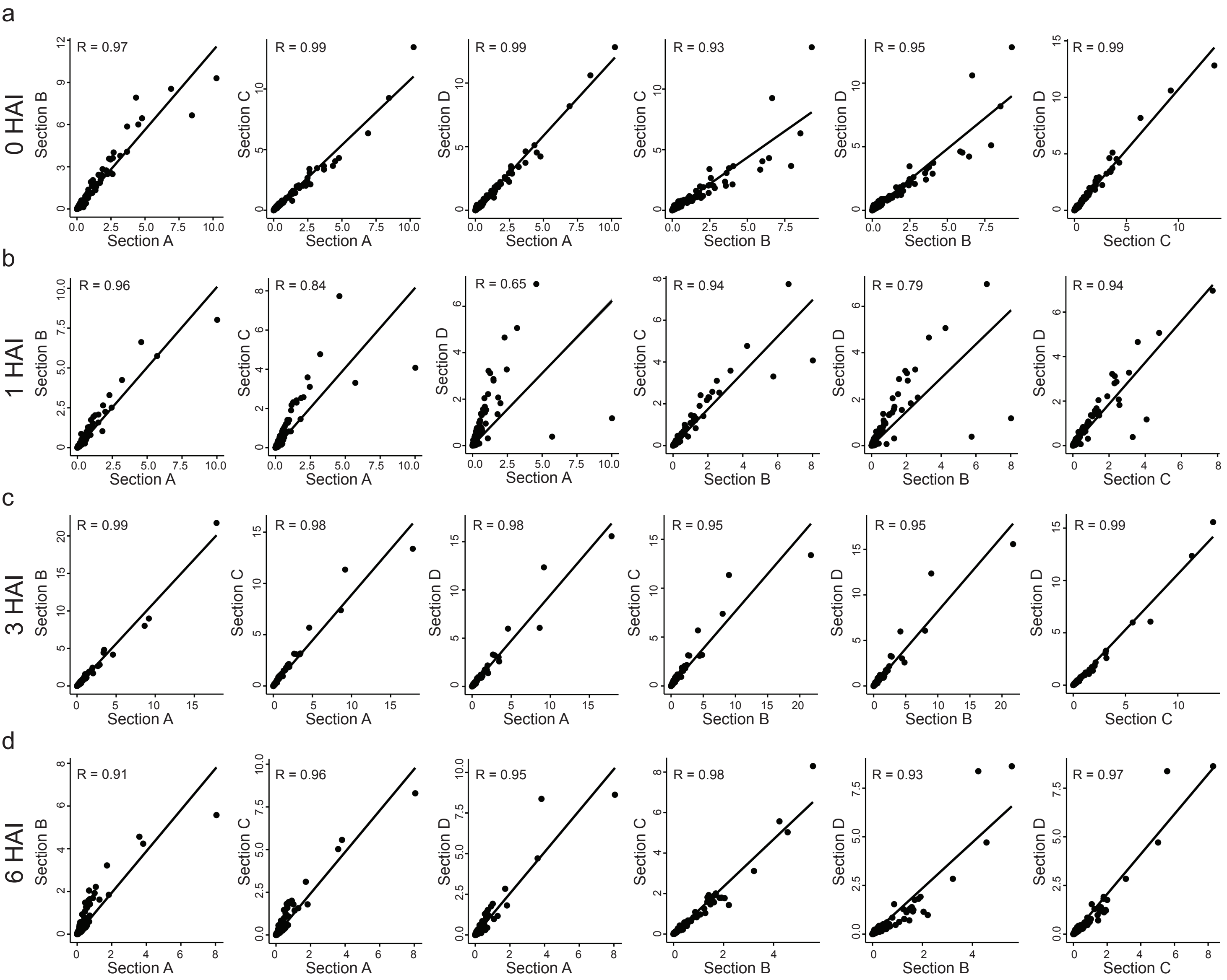


**Supplementary Figure 11.** Correlation between different sections at each time point.

### Supp Fig 13

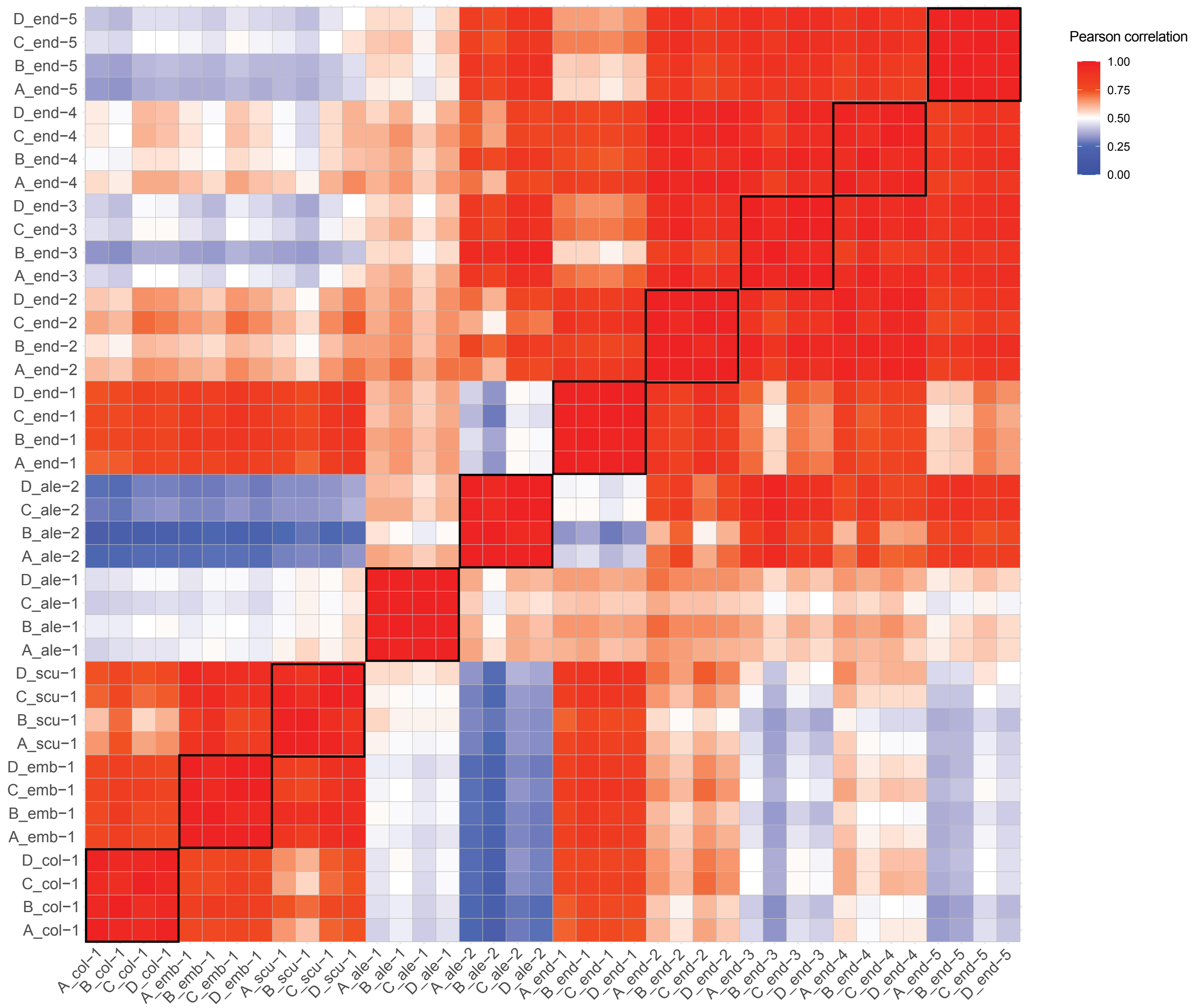


**Supplementary Figure 13.** Cluster correlation across different sections of 0 HAI.

### Supp Fig 14

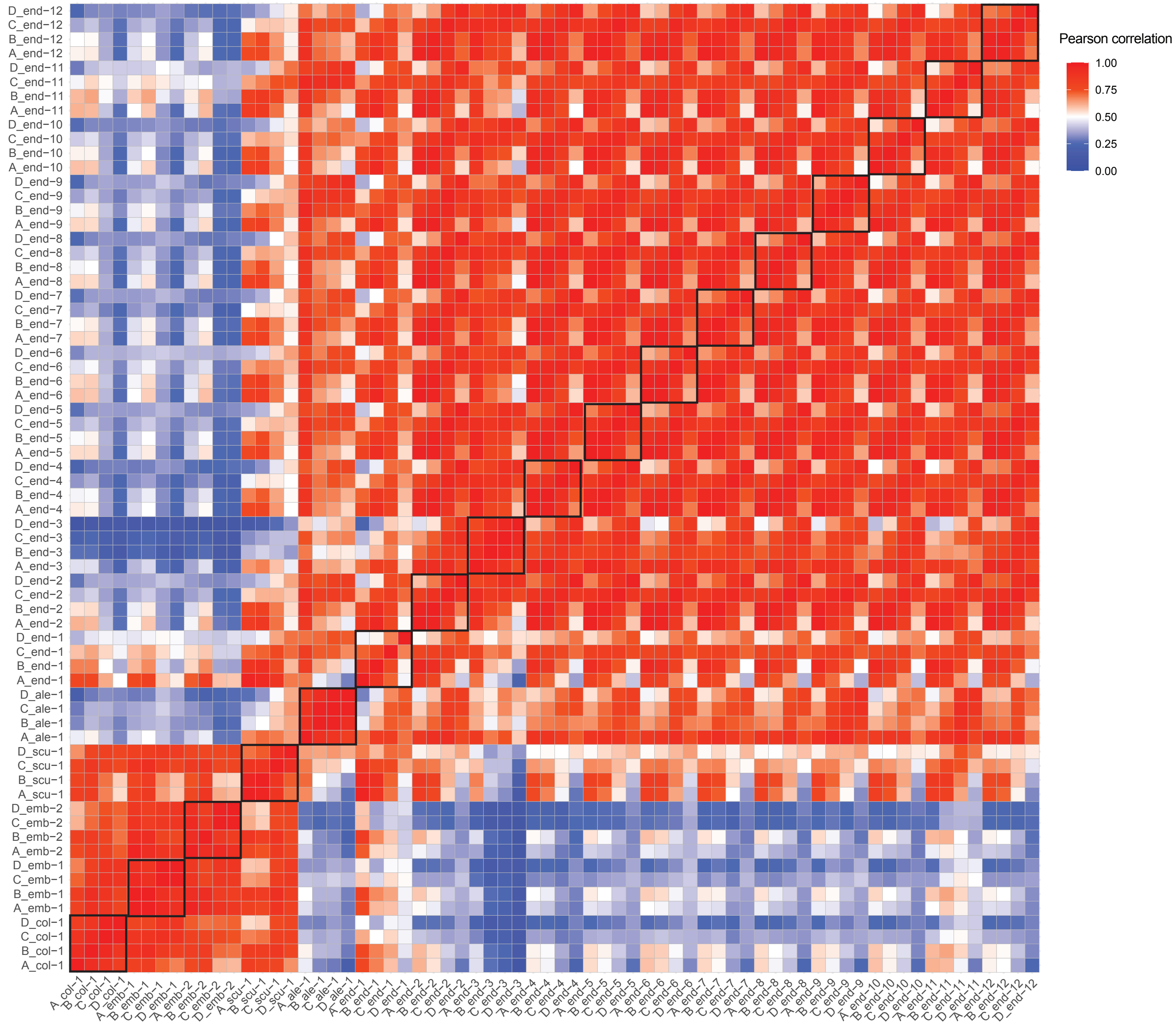


**Supplementary Figure 14.** Cluster correlation across different sections of 1 HAI.

### Supp Fig 15

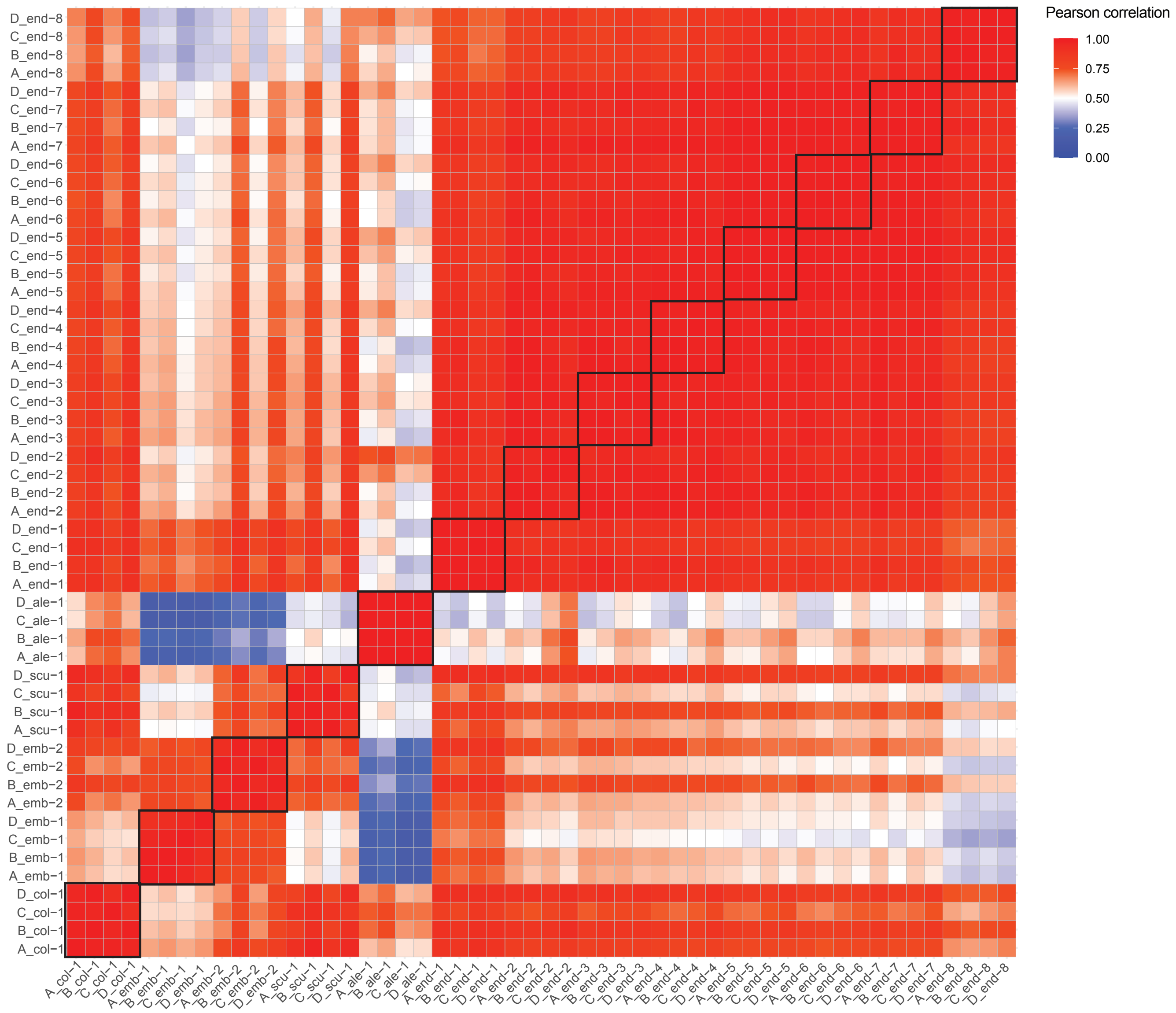


**Supplementary Figure 15.** Cluster correlation across different sections of 3 HAI.

### Supp Fig 16

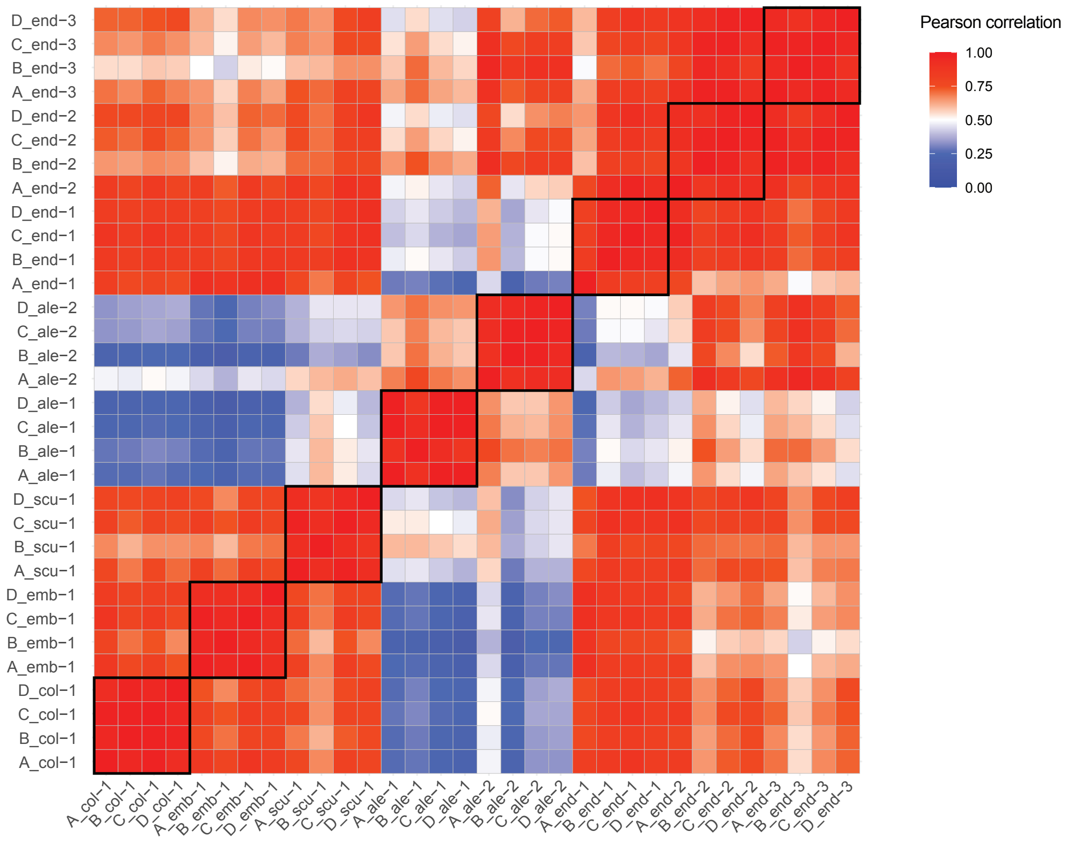


**Supplementary Figure 16.** Cluster correlation across different sections of 6 HAI.

### Supp Fig 17

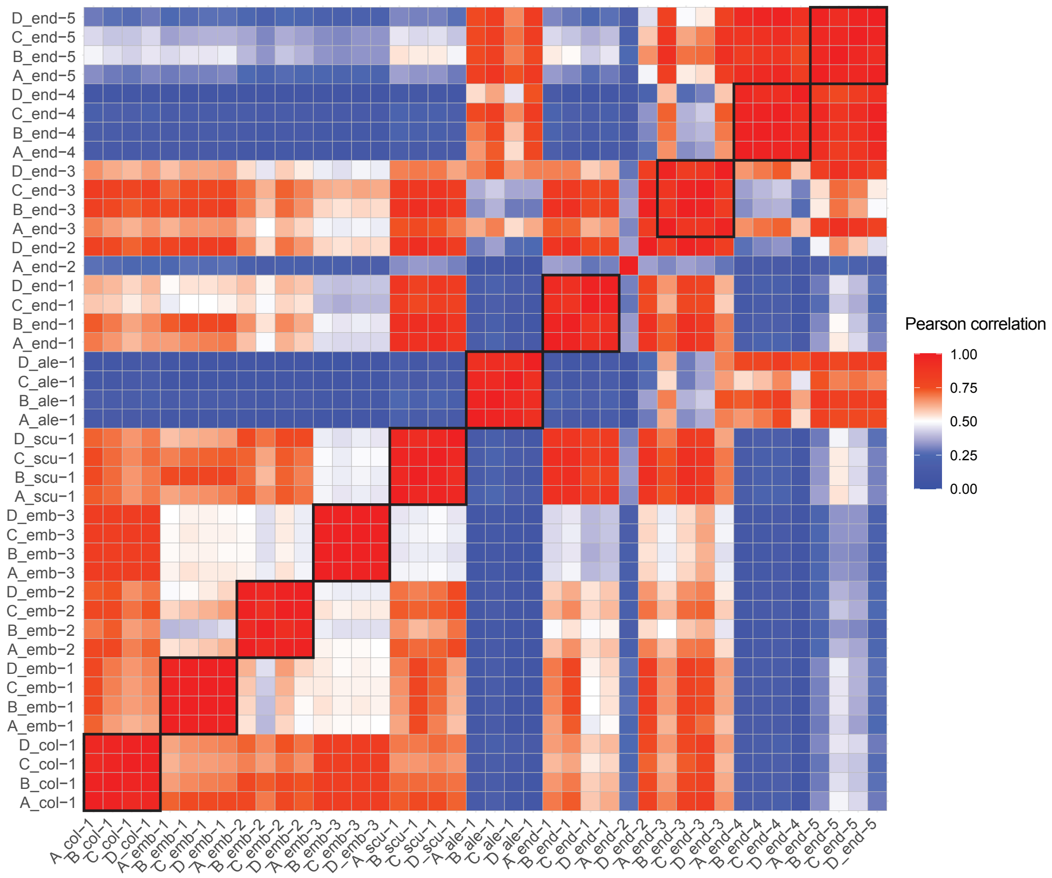


**Supplementary Figure 17.** Cluster correlation across different sections of 24 HAI.

### Supp Fig 19

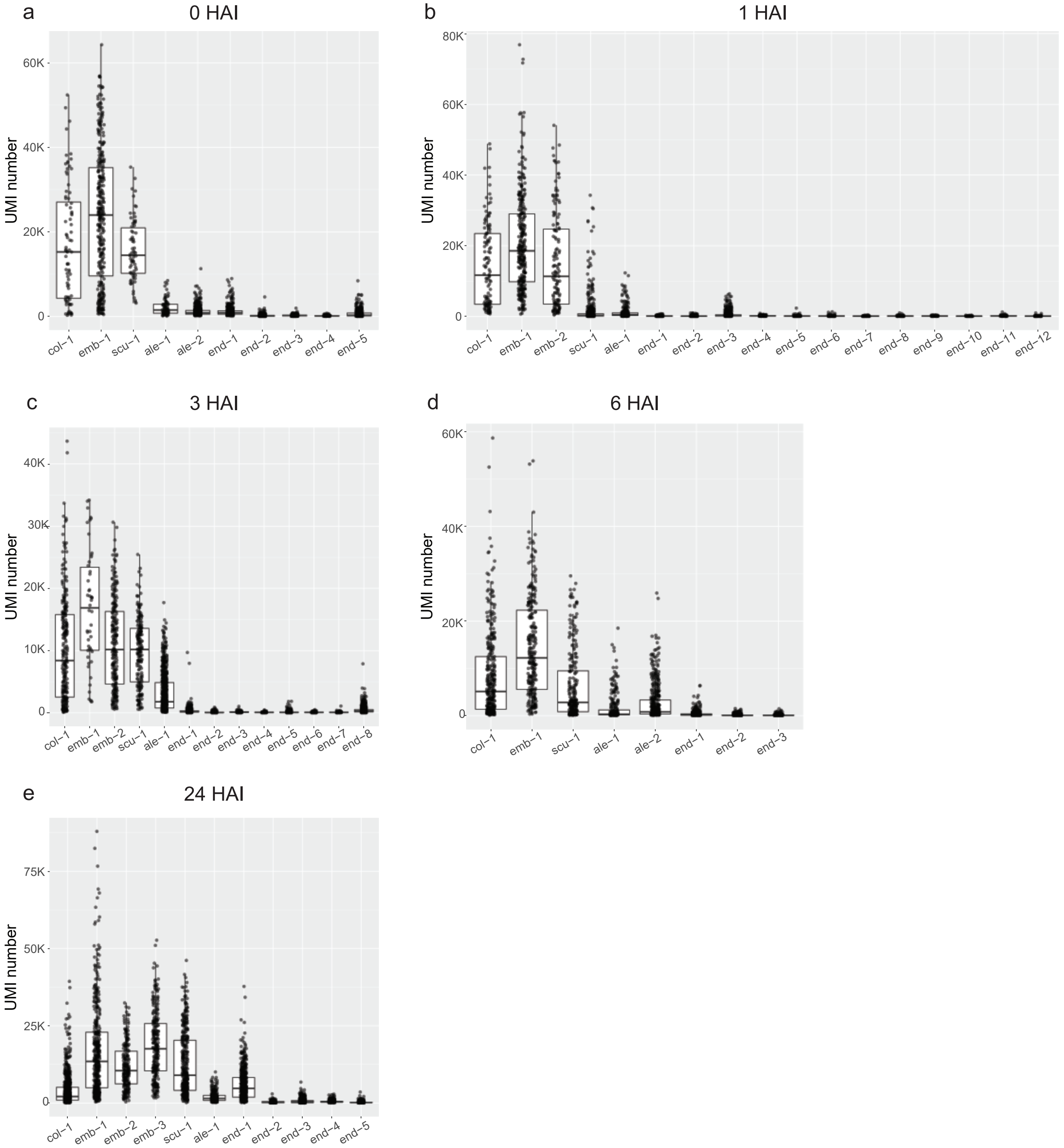


**Supplementary Figure 19.** UMI number of different clusters across different time points.

### Supp Fig 20

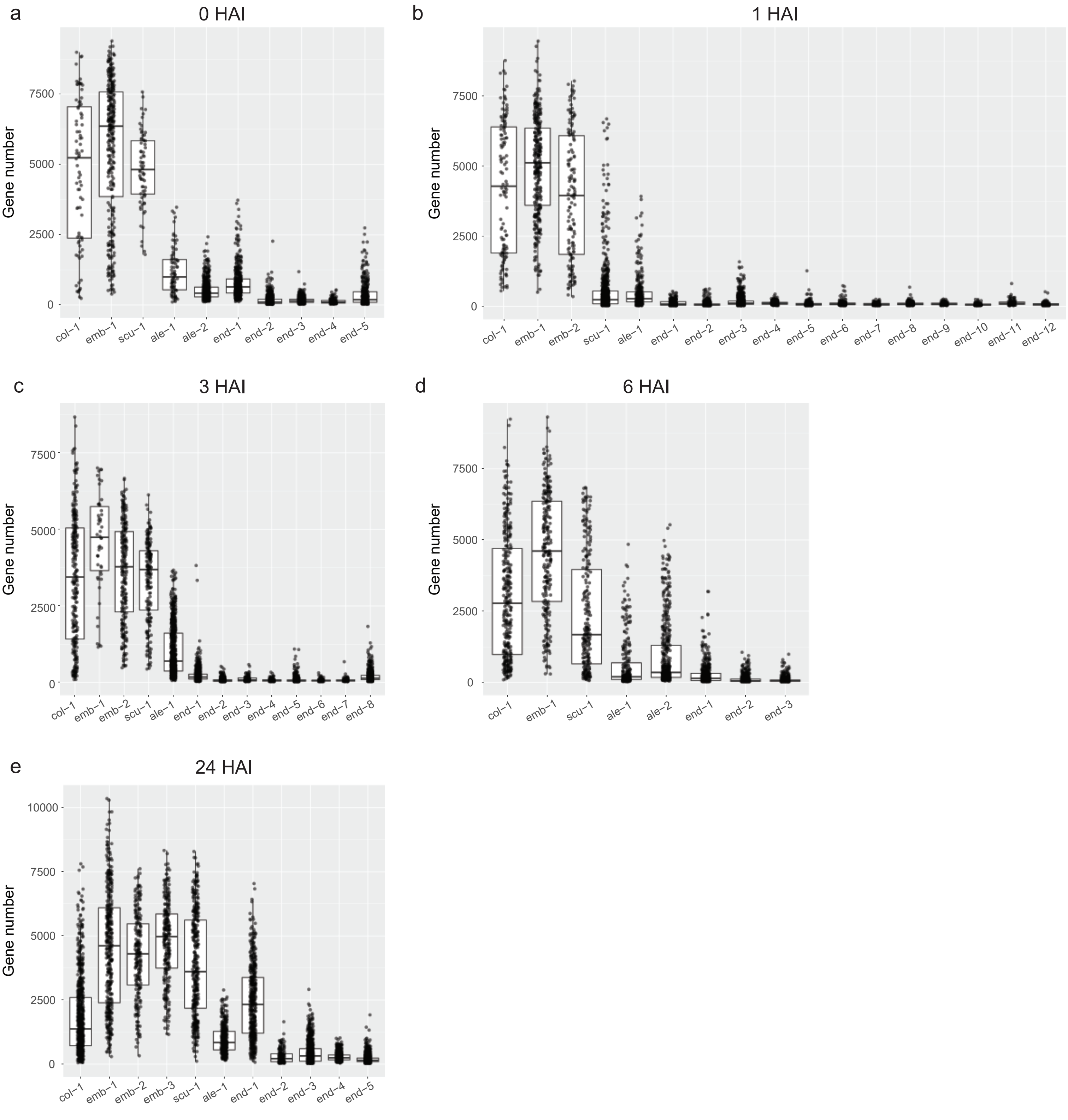


**Supplementary Figure 20.** Gene number of different clusters across different time points.

### Supp Fig 26

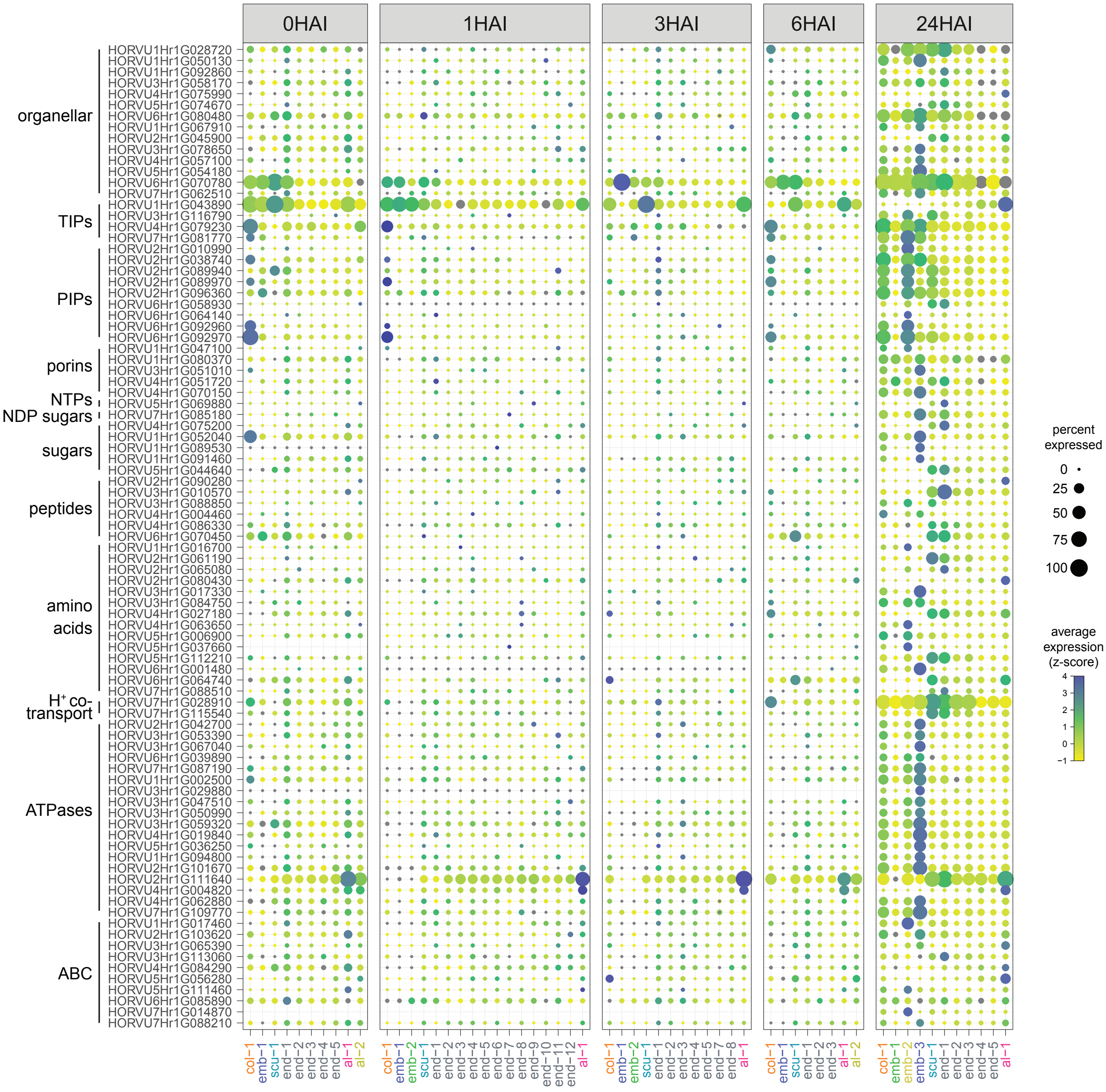


**Supplementary Figure 26.** Transport-related gene expression during barley germination.

### Supp Fig 27

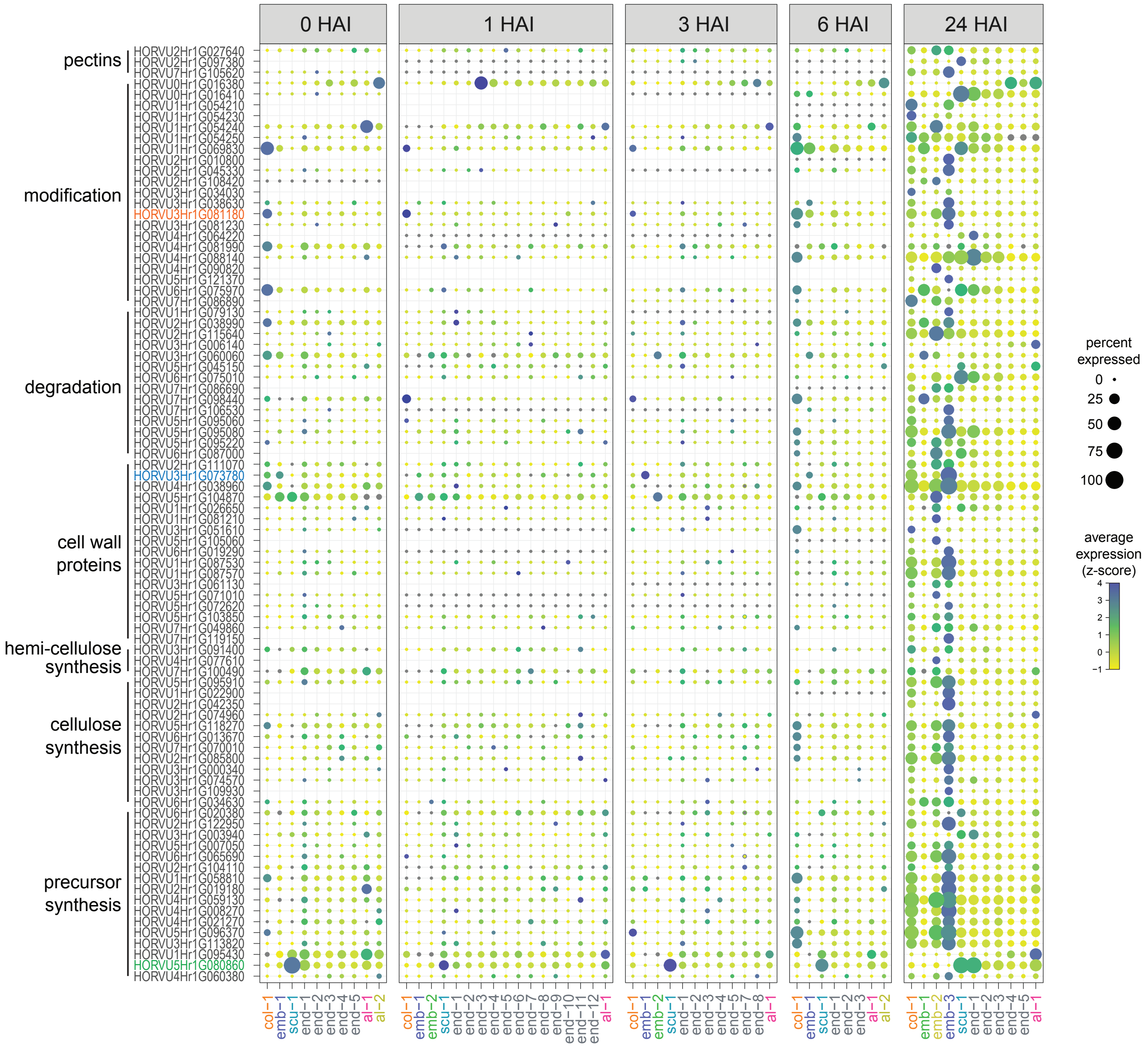


**Supplementary Figure 27.** Cell wall-related gene expression during barley germination.

### Supp Fig 28

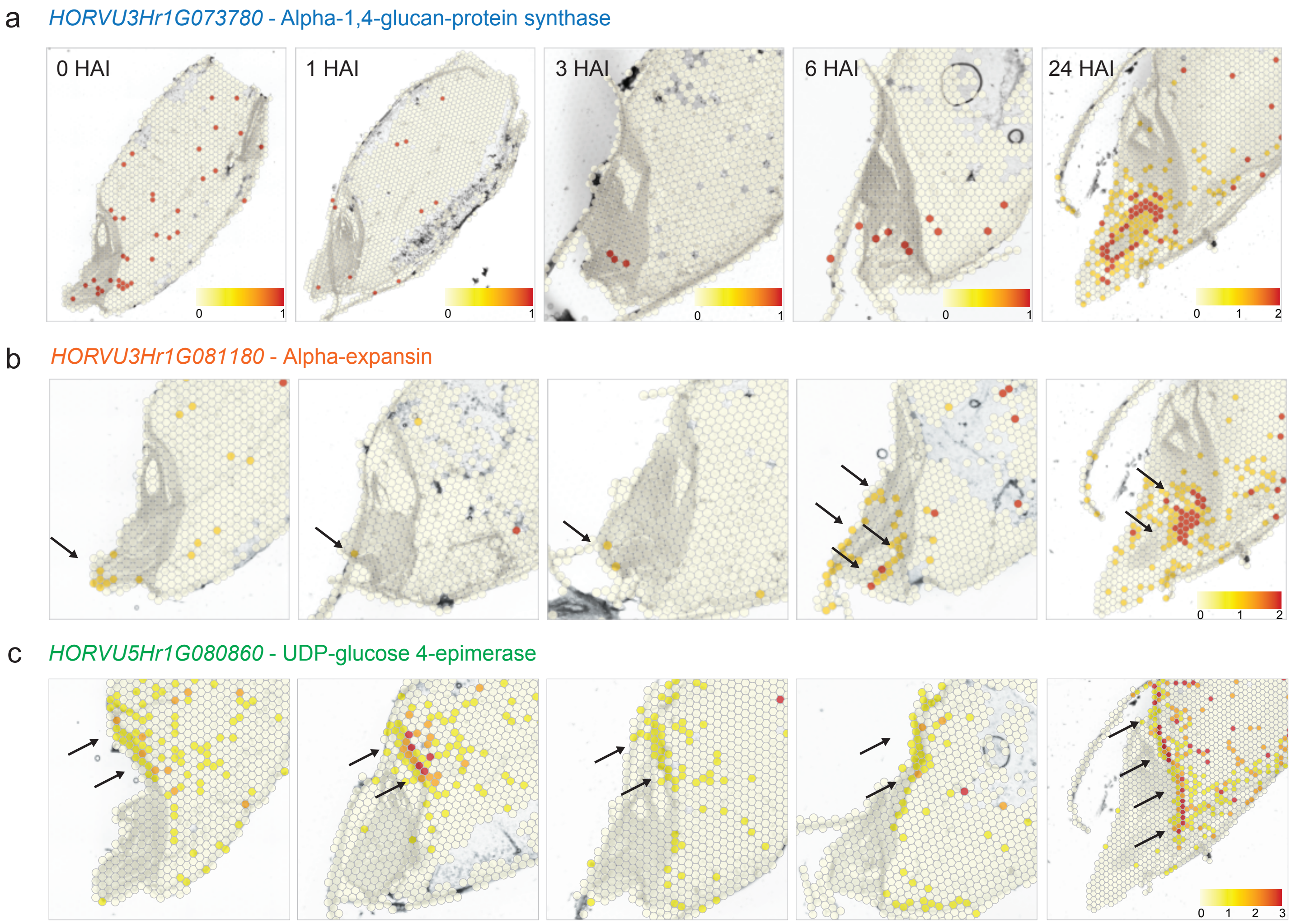


**Supplementary Figure 28.** Cell wall protein-related gene expression during barley germination.
