## Supplementary material for "Spatially Resolved Transcriptomic Analysis of the Germinating Barley Grain": Supp Fig 12

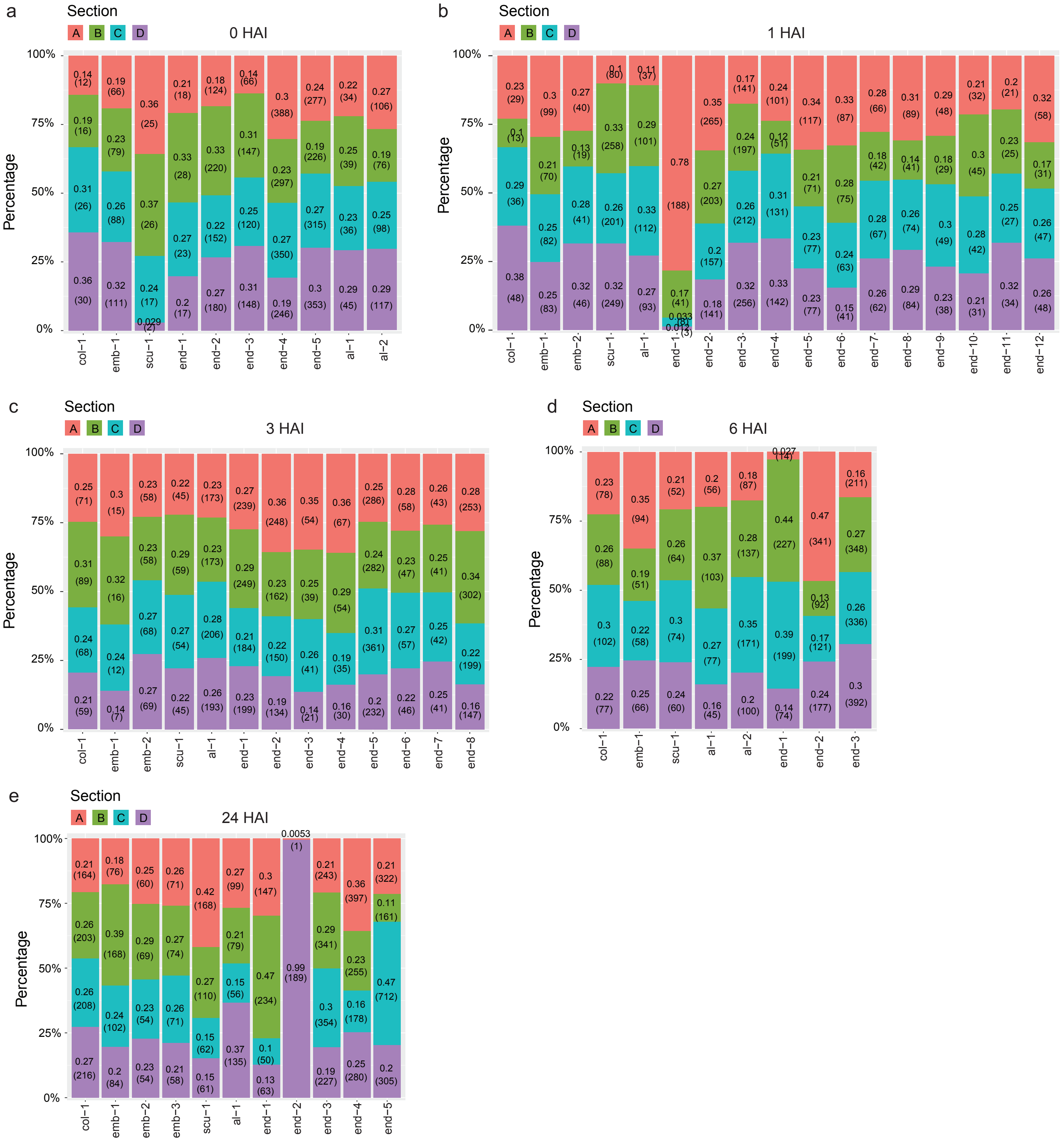


**Supplementary Figure 12.** Percentage of spots of four barley sections for each cluster for different time point.
