## Supplementary material for "Spatially Resolved Transcriptomic Analysis of the Germinating Barley Grain": Supp Fig 18

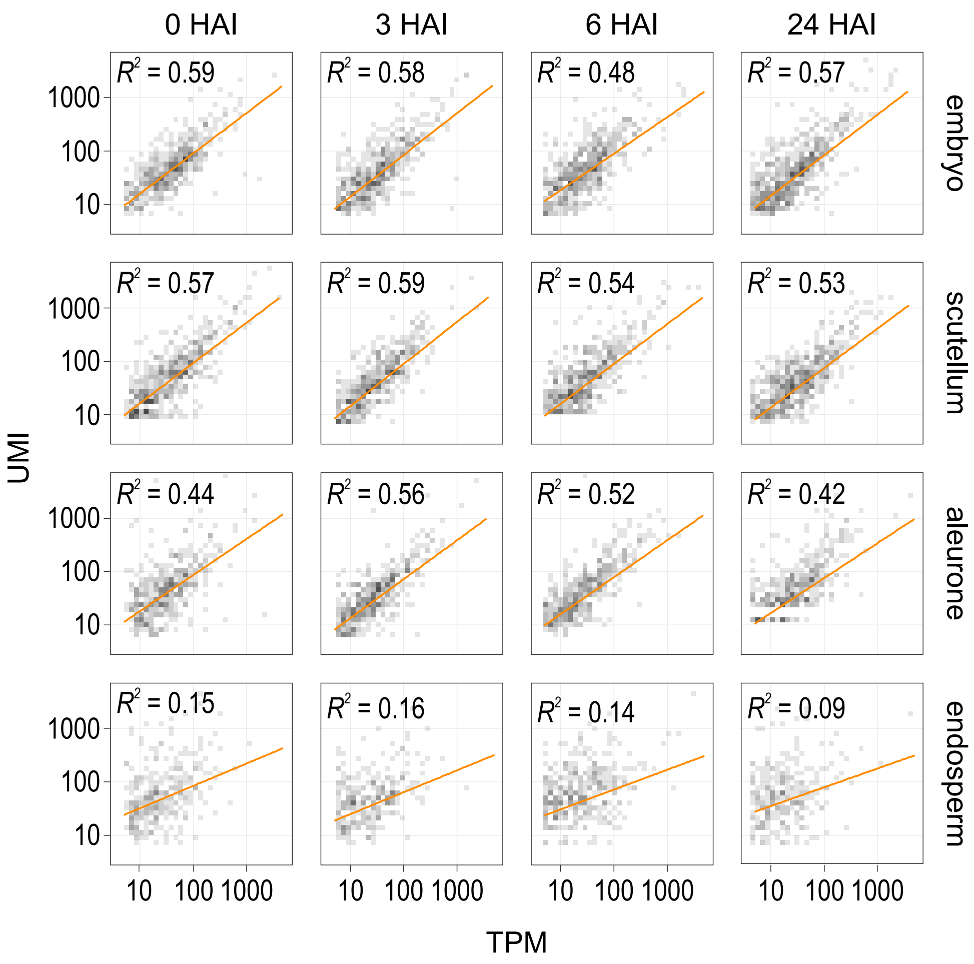


**Supplementary Figure 18.** Correlation of gene expression in STomics and tissue-specific RNA-seq experiments.
