## Supplementary material for "Spatially Resolved Transcriptomic Analysis of the Germinating Barley Grain": Supp Fig 21

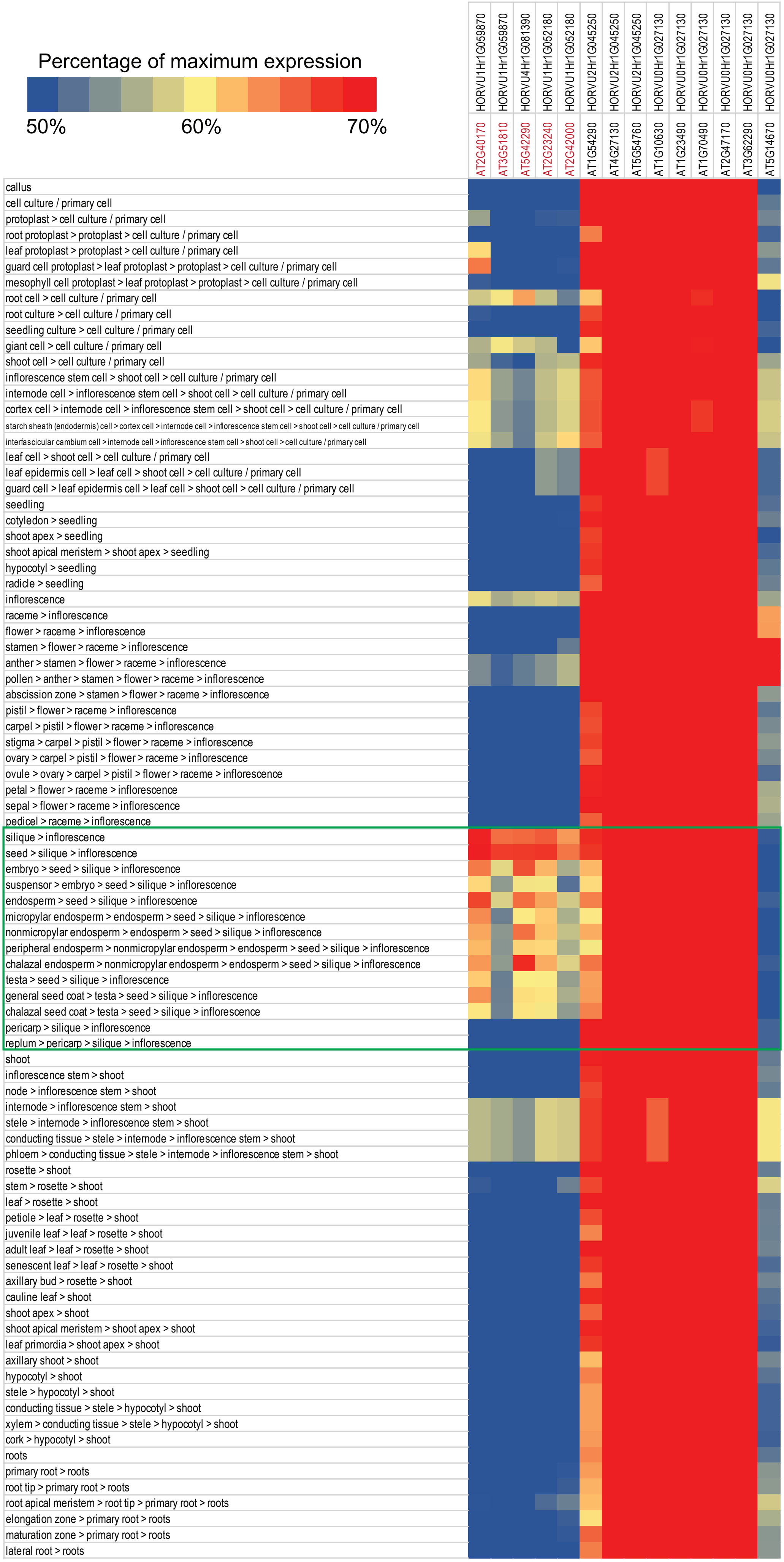


**Supplementary Figure 21**. Analysis of the tissue expression pattern of the Arabidopsis thaliana orthologs of the top 10 marker genes per cluster for endosperm at each time point.
