## Supplementary material for "Spatially Resolved Transcriptomic Analysis of the Germinating Barley Grain": Supp Fig 22

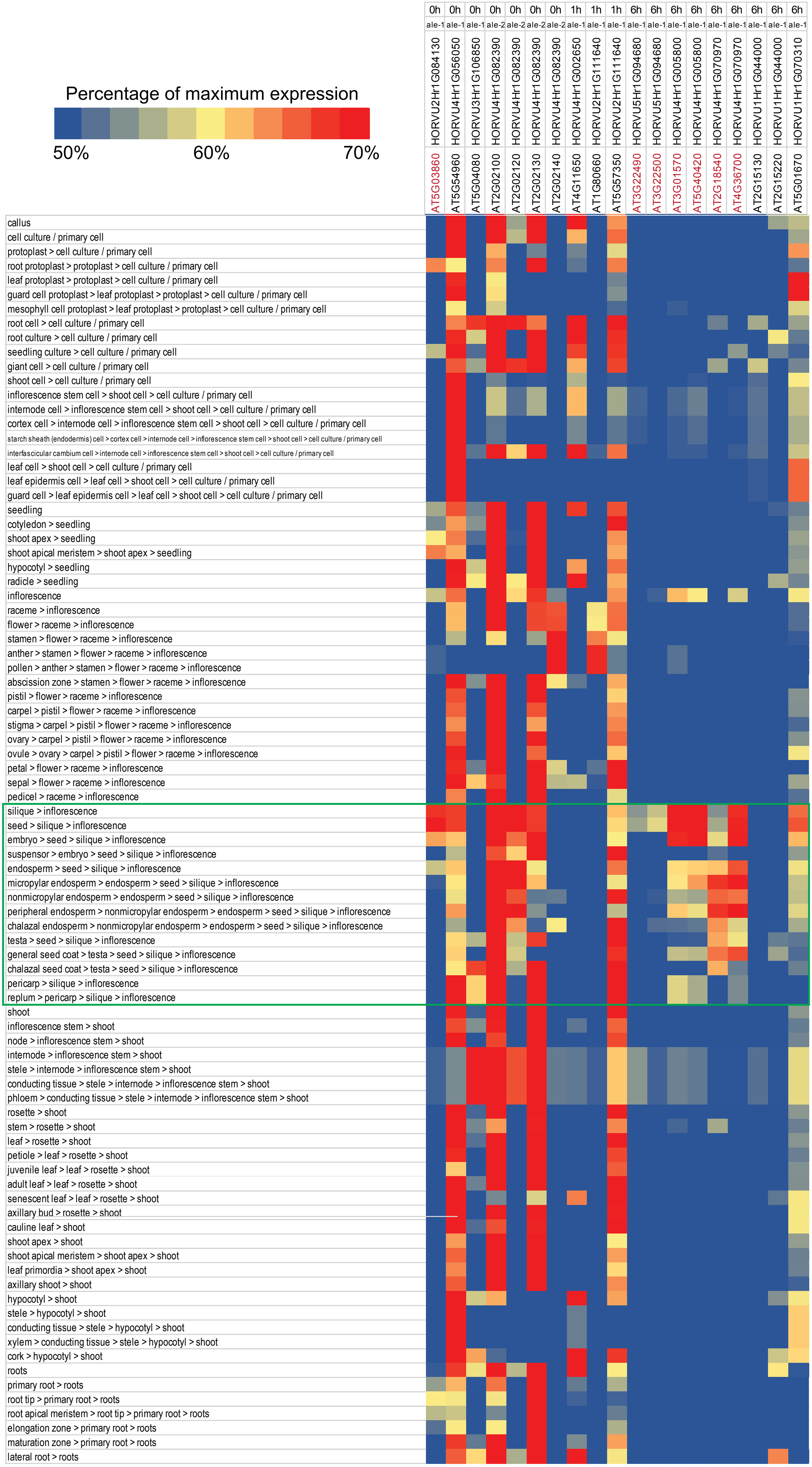


**Supplementary Figure 22.** Analysis of the tissue expression pattern of the Arabidopsis thaliana orthologs of the top 10 marker genes per cluster for aleurone at each time point.
