## Supplementary material for "Spatially Resolved Transcriptomic Analysis of the Germinating Barley Grain": Supp Fig 23

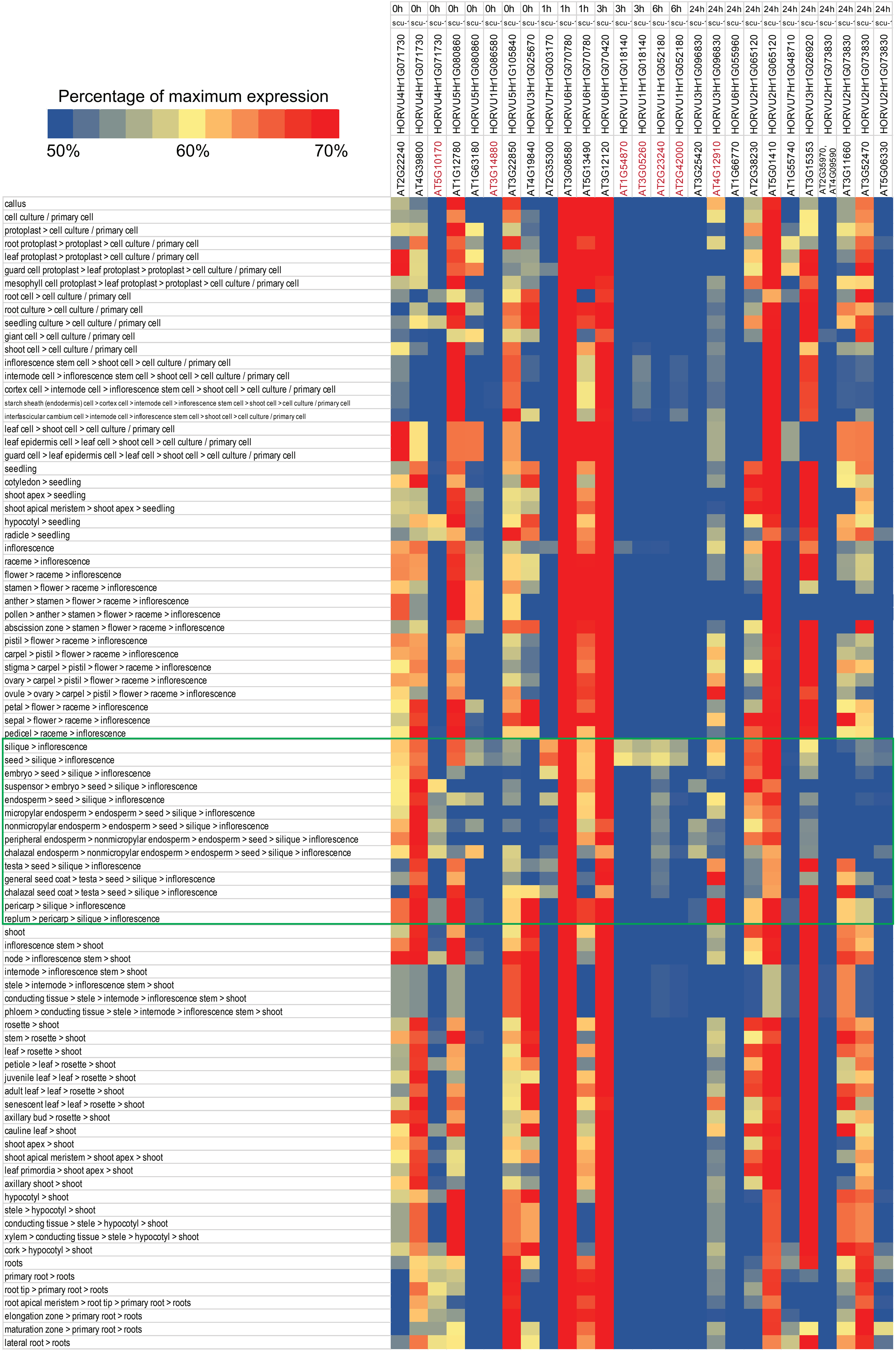


**Supplementary Figure 23.** Analysis of the tissue expression pattern of the Arabidopsis thaliana orthologs of the top 10 marker genes per cluster for scutellum at each time point.
