## Supplementary material for "Spatially Resolved Transcriptomic Analysis of the Germinating Barley Grain": Supp Fig 25

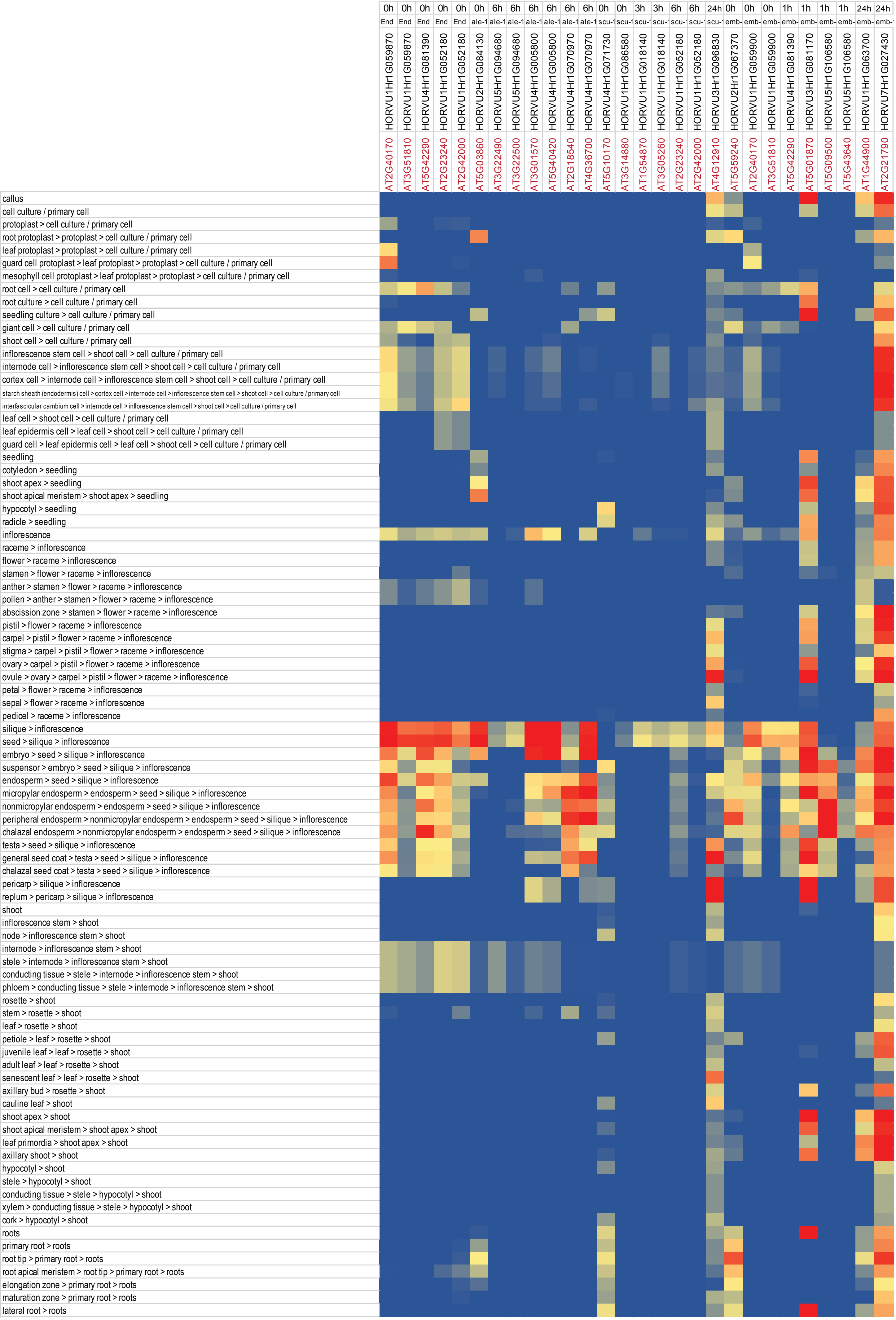


**Supplementary Figure 25.** Summary of the orthologous genes from Arabidopsis and barley that show similar tissue specific expression.
