## Supplementary material for "Spatially Resolved Transcriptomic Analysis of the Germinating Barley Grain": Supp Fig 29

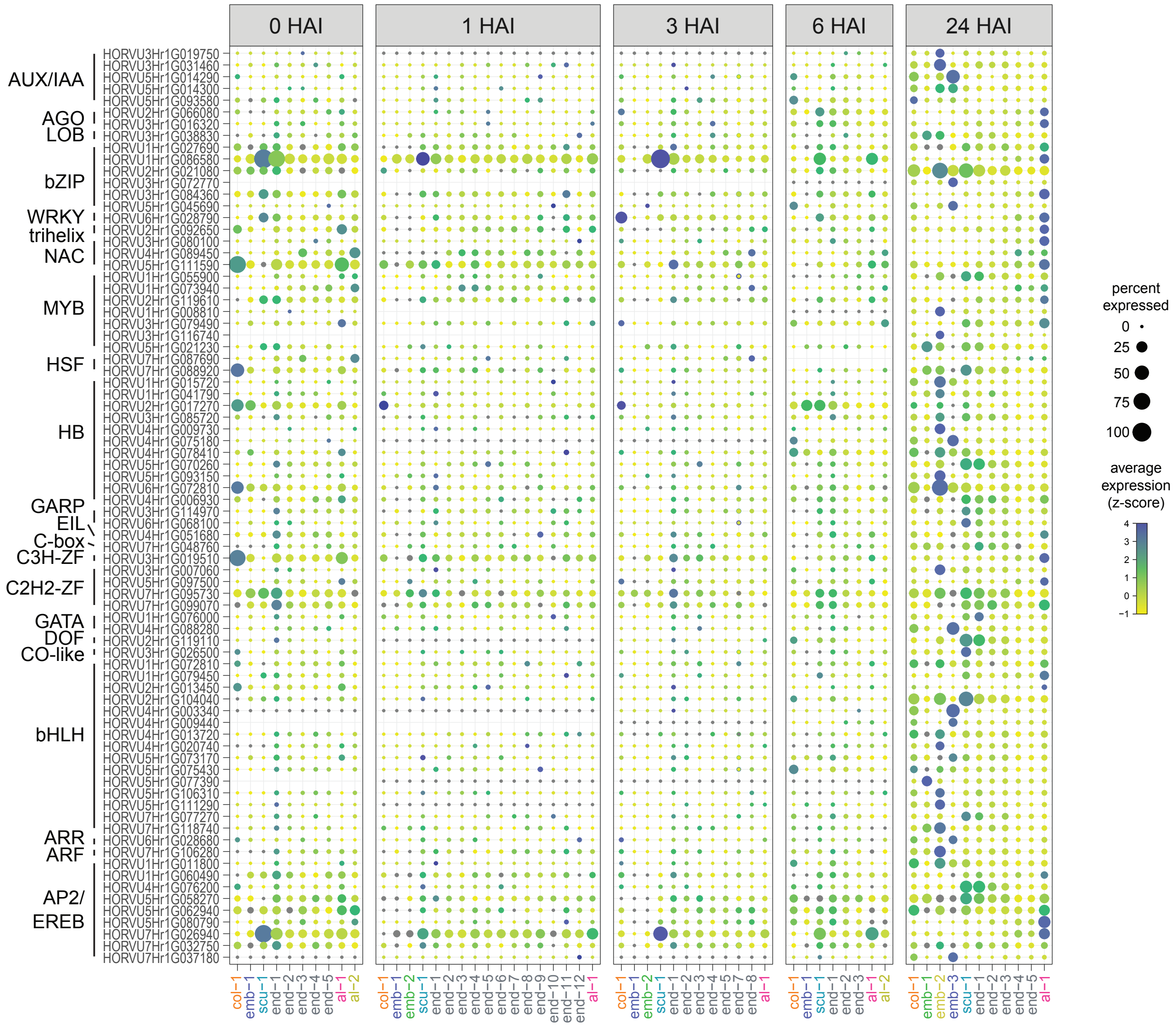


**Supplementary Figure 29.** Transcription factor-related gene expression during barley germination.
