## Supplementary material for "Spatially Resolved Transcriptomic Analysis of the Germinating Barley Grain": Supp Fig 30

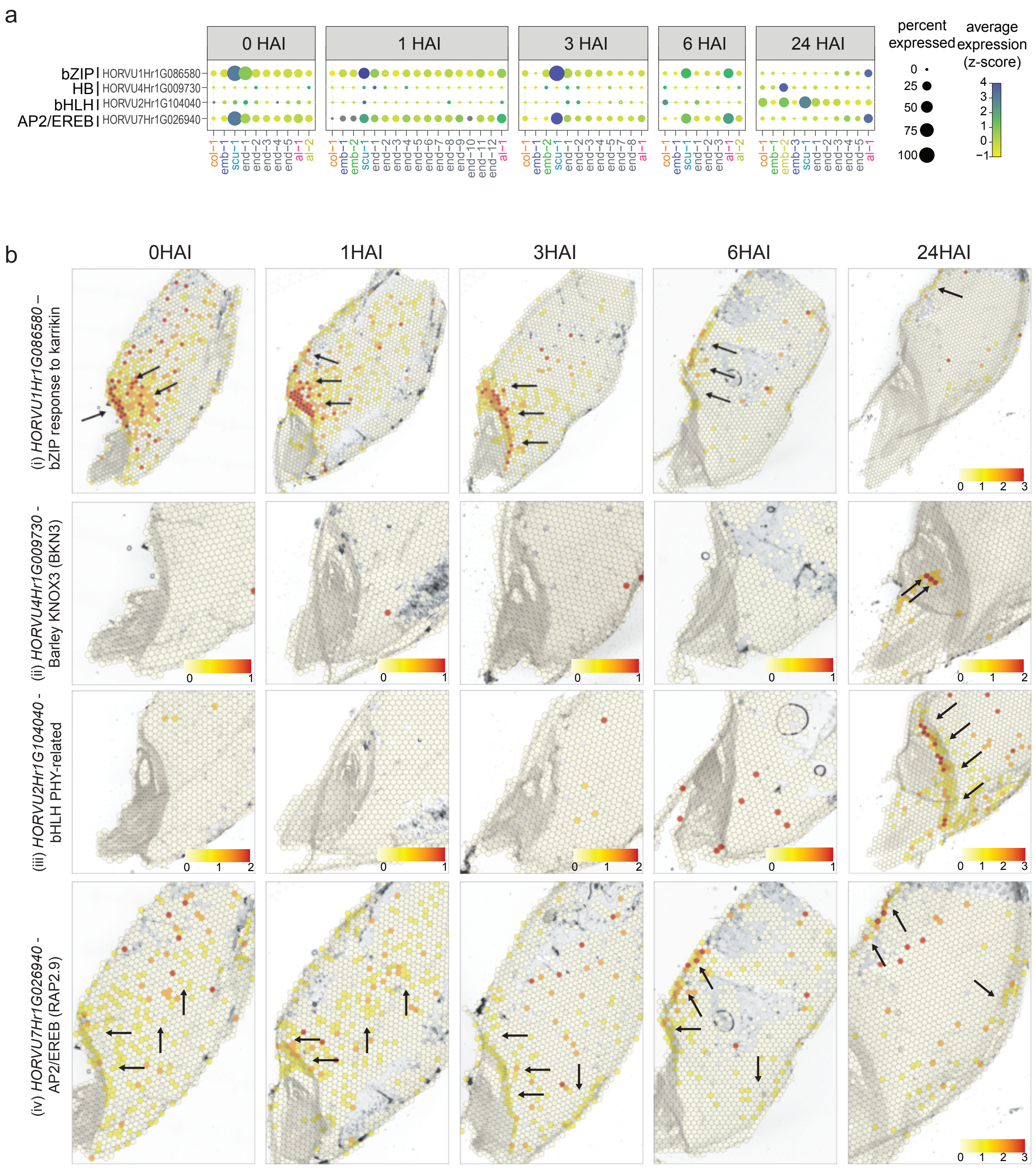


**Supplementary Figure 30.** Spatio-temporal expression of transcription factor-related genes during barley germination.
