## Supplementary material for "Spatially Resolved Transcriptomic Analysis of the Germinating Barley Grain": Supp Fig 31

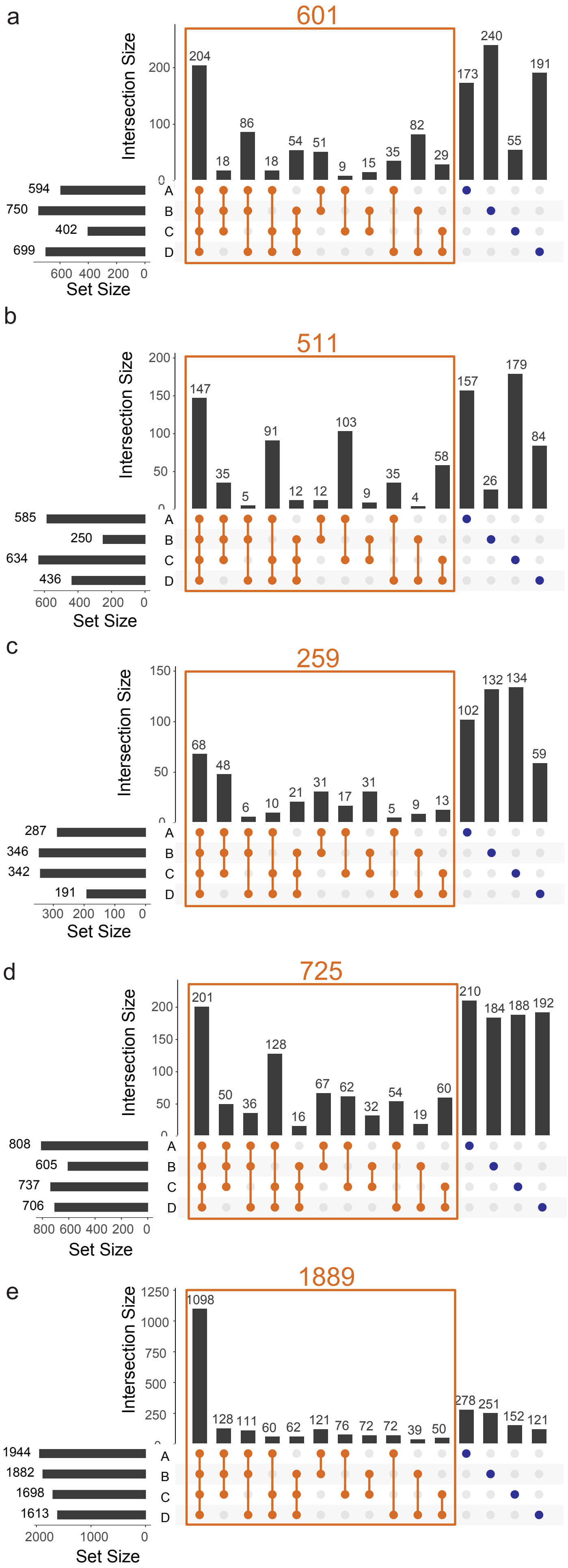


**Supplementary Figure 31.** Overlap of number of SVGs in different sections across different time points.
